# Mapping SOX9 transcriptional dynamics during multi-lineage differentiation of human mesenchymal stem cells

**DOI:** 10.1101/2021.11.29.470107

**Authors:** Kannan Govindaraj, Sakshi Khurana, Marcel Karperien, Janine N. Post

## Abstract

The master transcription factor SOX9 is a key player during chondrocyte differentiation, cartilage development, homeostasis and disease. Modulation of SOX9 and its target gene expression is essential during chondrogenic, osteogenic and adipogenic differentiation of human mesenchymal stem cells (hMSCs). However, lack of sufficient knowledge about the signaling interplay during differentiation remains one of the main reasons preventing successful application of hMSCs in regenerative medicine. We previously showed that Transcription Factor – Fluorescence Recovery After Photobleaching (TF-FRAP) can be used to study SOX9 dynamics at the single cell level. We showed that changes in SOX9 dynamics are linked to its transcriptional activity. Here, we investigated SOX9 dynamics during differentiation of hMSCs into the chondrogenic, osteogenic and adipogenic lineages. We show that there are clusters of cells in hMSCs with distinct SOX9 dynamics, indicating that there are a number of subpopulations present in the heterogeneous hMSCs. SOX9 dynamics data at the single cell resolution revealed novel insights about its activity in these subpopulations (cell types). In addition, the response of SOX9 to differentiation stimuli varied in these subpopulations. Moreover, we identified donor specific differences in the number of cells per cluster in undifferentiated hMSCs, and this correlated to their differentiation potential.

## Introduction

hMSCs are of great interest for regenerative therapy. Their potential to differentiate into chondrocytes, osteoblasts, and adipocytes substantiate therapeutic applications. hMSCs can be isolated from patients, can be differentiated *in vitro* and re-implanted in the patient body. Despite promising advances, clinical applications are largely unsuccessful due to several reasons, including sub-optimal differentiation and lack of sufficient knowledge about the signaling interplay, which is necessary to modulate the differentiation processes [1, 2].

Master transcription factors play a key role in orchestrating signaling cascade during differentiation. For example, SOX9 [3], RUNX2 [4] and PPARγ [5] are tightly regulated by the signaling interplay during chondrogenic, osteogenic and adipogenic differentiation of hMSCs, respectively. Although much is known about their collective mRNA and protein expression levels during various stages of differentiation, little is known about the real-time dynamics of these transcription factors [2]. Mapping real-time dynamics will help us to understand the response of transcription factors to changes in signaling network activities during the differentiation process. Given the complex hMSC heterogeneity within a donor and the large cell-cell variability, static gene and protein quantification methods do not yield the much-needed dynamic information about protein activity.

SOX9 is the master transcription factor for chondrocyte differentiation, cartilage development and homeostasis. SOX9 also plays a role in endochondral ossification and formation of growth plate [6]. In addition, SOX9 is involved in the development of testis and its mis-regulation is linked to various types of cancers including breast, prostate and chondrosarcoma [7]. Even a small change in SOX9 function during development will have profound impact in the function of skeletal system. Mutations in the SOX9 gene are known to cause campomelic dysplasia, a disease characteristic of cartilage and bone malformation [8]. SOX9 expression is increased during mesenchymal condensation and early chondrogenesis. SOX9 directly activates key chondrocyte genes aggrecan and type II collagen. Its expression is reduced or turned off during differentiation into other lineages [7].

In contrast, reduced SOX9 expression is linked to loss of cartilage homeostasis, incomplete MSC differentiation, and hypertrophy, etc. Therefore, others investigated methods for more efficient differentiation based on overexpression of SOX9 [9, 10]. Furthermore, SOX9 and its target gene expression levels are known to fluctuate during differentiation of hMSCs and are modulated in various differentiation lineages [11, 12]. However, signaling and transcriptional regulation of SOX9 and its target genes are not yet fully understood.

We have previously shown that the average TF-FRAP rates correlate to DNA binding (ChIP assay) and transcriptional activity (gene expression analysis) [13]. However, these bulk measurements did not give sufficient information about SOX9 transcriptional activity in a time-resolved manner at the subpopulation level in a heterogenic cell population in, for example, MSCs.

Increased temporal resolution of SOX9 transcriptional activity will help us to understand more about the signaling interplay during differentiation. We applied our recently developed TF-FRAP to study the SOX9 dynamics in proliferating and differentiating hMSCs isolated from bone marrow. To understand SOX9 transcriptional activity during differentiation, we differentiated hMSCs into chondrogenic, osteogenic, and adipogenic lineage and measured SOX9 dynamics at day 0, 2, 8 and 15 or 23. TF-FRAP captured SOX9 dynamics with higher spatiotemporal resolution. Interestingly, our TF-FRAP revealed how SOX9 dynamics changed in various subpopulations of hMSCs during differentiation per lineage and time point.

## Materials and Methods

### Vector-plasmids

The vector expressing SOX9-mGFP was constructed by cloning mGFP (PS100040, Origene) with the C-terminal of wild type SOX9 (RC208944, Origene) using SgfI and MluI restriction sites. The correct reading frame of the fusion construct was verified by sequencing and its functionality was confirmed by mutation studies [13].

### Cell Culture and Transfection

hMSCs were isolated from bone marrow from patients with no known musculoskeletal diseases. hMSCs were cultured in αMEM (22571-038, Gibco), supplemented with 10% FBS (F7524, Sigma), 200mM glutaMax (35050-38, gibco), 20 mM ascorbic acid 2 phosphate (AsAP, Sigma, A8960), 100 ng/ml bFGF (Neuromics, RP80001), and 100 U/ml of Penicillin-Streptomycin (15140122, gibco) at 37 ⁰C with 5% CO_2_. hMSCs were expanded and used within 4 passages.

C20/A4 cells were cultured in DMEM (Gibco, USA) with 10% FBS.

Human primary chondrocytes were cultured in chondrocyte proliferation media containing DMEM supplemented with 10% FBS, 20 mM ascorbic acid 2 phosphate (AsAP, Sigma, A8960), L-Proline (40 μg/ml) and non-essential amino acids, at 37 ⁰C with 5% CO_2_.

For transfection, cells were seeded on a sterile glass coverslip (40,000 per well) placed inside the well. Lipofectamine 3000 with P3000 Reagent (Life Technologies, USA) was used for transfection of hMSCs and Lipofectamine LTX with Plus reagent was used for transfection of C20/A4 cells and hPCs and the manufacturer’s protocol was followed.

### Cell synchronization

hMSCs were seeded on microscopic coverslip placed inside a 24-well plate in hMSC proliferation media. Next day, cells were transfected as described above and maintained in proliferation media with serum for about 8 hours to allow cells to produce SOX9-mGFP protein. For TF-FRAP: Transfected cells were maintained in serum free proliferation media 24 hours prior to TF-FRAP measurements. During TF-FRAP measurements, cells were maintained in the imaging buffer as described below. For imaging SOX9-mGFP nuclear localization pattern: for 24+0h time point, transfected cells were maintained in serum free proliferation media 24 hours prior to imaging. For 24+6h time point, post 24h of serum starvation, cell cycle was again started by replacing the media with proliferation media with serum. Cells were imaged 6h after the start of cell cycle, so the time point 24+6h. Cells in two different coverslips were used for 24+0h and 24h+6h time points study. During imaging, cells were maintained in the imaging buffer, at 37⁰C.

### hMSCs differentiation

hMSCs were differentiated into chondrogenic, osteogenic and adipogenic lineages by culturing them in their respective differentiation media from day 0. Culture media was refreshed every 3 – 4 days. Chondrogenic differentiation medium: hMSCs (10,000 cells/cm^2^) were cultured in DMEM (Gibco) supplemented with 1% of Insulin-Transferrin-Selenium (ITS) mix (41400045, Gibco), 40 μg/ml of L-proline, 50 μg/ml of AsAP, 1% of sodium pyruvate (S8636, Sigma), 100 U/ml of Penicillin-Streptomycin, and freshly added TGFβ1 (10 ng/ml, 7754-BH, R&D Systems) and 10^-7^ M dexamethasone (Dex, D8893, Sigma). Osteogenic differentiation medium: hMSCs (1,000 cells/cm^2^) were cultured in αMEM supplemented with 10% FBS, 100 U/ml of Penicillin-Streptomycin, 200 mM GlutaMax, 50 μg/ml AsAP and freshly added Dex (10^-8^ M). Adipogenic differentiation medium: hMSCs (10,000 cells/cm^2^) were cultured in αMEM supplemented with 10% FBS, 100 U/ml of Penicillin-Streptomycin, 200 mM glutaMax, 50 μg/ml AsAP, freshly added Dex (10^-6^ M), 10 μg/ml Insulin (I9278, Sigma), 0.5 mM IBMX (I5879, Sigma) and 0.2 mM Indomethacin (57413, Sigma).

### Imaging Buffer

Imaging was performed in Tyrode’s buffer with freshly added 20 mM glucose (Gibco) and 0.1% BSA (Sigma) [14]. Tyrode’s buffer is composed of 135 mM NaCl (Sigma), 10 mM KCl (Sigma), 0.4 mM MgCl2 (Sigma), 1 mM CaCl_2_ (Sigma), 10 mM HEPES (Acros organics), pH adjusted to 7.2, filter sterilized (0.2 μm) and stored at -20⁰C.

### Transcription factor-Fluorescence Recovery After Photobleaching

Proliferating or differentiating hMSCs were plated on poly-l-lysine (0.01%, Sigma), coated glass cover slips 2 days before transfection and were transiently transfected with SOX9-mGFP a day before of FRAP experiments. For chondrogenic differentiating cells, FBS was added to the chondrogenic medium for attachment of the cells to the coverslip. FRAP was performed at four time points, i.e., 0 day (proliferation), day 2, day 8 and day 15 or 23 for all the three lineages. Cells were maintained in imaging buffer during FRAP measurements and the mobility of transcription factors was measured at the steady-state (i.e. without other stimuli or FBS added). FRAP measurements were performed using a Nikon A1 laser scanning confocal microscope (Nikon, Japan) with a 60X/1.2 NA water immersion objective, 488 nm Argon laser at 0.35% (0.12 μW at the objective) laser power for SOX9-mGFP. The temperature was maintained at 37 ⁰C with an OkaLab temperature controller. A frame size of 256x256 pixels, covering the whole nucleus, was scanned. The pixel size was 0.12 μm. A representative circular region of 2.9 μm diameter was bleached with one iteration (60 ms) of 50% (34.3 μW) laser power. Twenty-five pre-bleach images were taken and the last 10 pre-bleach fluorescence intensity values were averaged to normalize the post-bleach fluorescence recovery curve. After bleaching, imaging was performed at 4 frames/sec for 60 sec post-bleach. FRAP experiments were performed on at least 40 cells per condition. To assess the statistical significance between the conditions Mann-Whitney U tests were applied using Origin® software. Matlab^TM^ was used to analyze the FRAP data and the script is available upon request. A diffusion uncoupled, two-component method was used to interpret our FRAP results as mentioned in [13].

### Clustering and statistical analysis

We applied unsupervised hierarchical clustering to cluster SOX9 dynamics data. TF-FRAP variables, such as Immobile Fraction (IF), Recovery half-time of A_1_ and A_2_ were used as input for cluster analysis. The following clustering parameters were used: Distance type: Euclidean, Cluster method: Furthest neighbor and Find Clustroid: Sum of distances. The distinct shapes of FRAP curves served as good references to identify and determine the number of clusters in our FRAP data. Clusters within a differentiation lineage at different time points that showed no significant difference (at P>=0.05, Mann-Whitney U-test) for at least two FRAP variables, were considered as same clusters, and were given the same letter/ identifier. OriginPro® (Origin Labs, USA) was used for clustering analysis.

### Generation of heatmaps

For convenience, clusters as segregated by hierarchical clustering were reordered per time-point within the differentiation lineage in ascending order of immobile fraction values. All three FRAP variables, IF, recovery half-time (t ½ ) of A_1_ and t ½ of A_2_ , were normalized between 0 and 100. Within a variable, the lowest and highest values were set to 0 and 100 respectively. OriginPro® was used to make heat-maps.

### Immunofluorescence

hMSCs cultured on microscopy coverslips were fixed for 10 min with freshly prepared 4% paraformaldehyde in PBS, pH7.2. Cells were washed with ice cold PBS, 3 times with 5 min interval. Cells were blocked and permeabilized for 15 mins with blocking solution containing 1 mg/ml BSA and 0.1% Triton X-100 in PBS. Cells were incubated with rabbit anti-human SOX9 primary antibody (AB5535, Millipore) in the blocking solution for 30 min at 37°C. Cells were washed with PBS, 3 times with 5 min intervals. Cells were incubated with goat anti-rabbit AF-568 antibody (ab175471, Abcam) in the blocking solution (with 0.05% Triton X-100) for 30 min at 37C. Cells were washed with PBS, 3 times with 5 min interval. Cell were stained with DAPI (1:100 dilution to 5 ng/ml) for 5 min in PBS. Cells were washed with PBS, 3 times with 5 min interval and mounted to microscopy glass side using Vecta shield mounting medium (company?). Cells were imaged using a Nikon A1 Confocal microscope with 60x water immersion objective with 1.2 NA.

### Histology Staining

Differentiation of hMSCs into chondrogenic, osteogenic and adipogenic lineage is followed by Alcian blue, ALP and Oil Red O staining, respectively, at various time points. Alcian blue staining was used to stain GAG produced by chondrogenic differentiated hMSCs at day 0, day 2, day 8 and day 23. Cells were fixed with 10% buffered formalin (HT501128, Sigma) for 15 mins and washed twice with ice cold PBS. Cells were incubated with freshly prepared Alcian Blue 8GX (A3157, Sigma) staining solution (0.5% w/v in 1M HCl, pH 1.0) for 30 mins and the cells were rinsed with dH_2_O until the staining solution is washed off. ALP staining was used to stain alkaline phosphatase activity by osteogenic differentiated hMSCs at day 0, day 2, day 8 and day 14. ALP staining kit (S5L2-1KT, Sigma) was used for the staining and the manufacturer’s protocol was followed. Oil Red O staining was used to stain neutral triglycerides and lipids produced by adipogenic differentiated hMSCs at day 0, day 2, day 8 and day 23. Cells were fixed with 10% buffered formalin for 15 mins and washed twice with ice cold PBS and dH_2_0. Cells were incubated in 60% isopropanol for 5 mins and then in freshly filtered Oil Red O staining solution for 5 mins and the cells were rinsed with dH_2_O until the staining solution was washed off (approximately 2 – 3 minutes). Cells were observed using a Nikon ECLIPSE TE300 microscope with 4x (0.13 NA) objective for all three staining.

### mRNA isolation and RT-qPCR

Chondrogenic, osteogenic and adipogenic differentiating and proliferating hMSCs were separately cultured in a 12-well plate for different time points. Proliferating hMSCs were considered as 0 hr control and for differentiating cells, gene expression pattern was studied at three time-points: day 2, day 8 and day 15 or 23 (for all 3 lineages). mRNA was isolated using TRIzol^TM^ (15596026, Invitrogen) and linear acrylamide as a co-precipitant, according to the manufacturer’s protocol. Purity and concentration of RNA samples were measured by Nanodrop 2000 (Thermo Fisher). cDNA was synthesized from total RNA with iScript cDNA synthesis kit (Bio-Rad). Real-time PCR analysis was carried out using SYBR Green mix (Bioline) in a Bio-Rad CFX-100 RT-qPCR. Gene expression is reported as the relative fold-change (ΔΔCt) [15] and is normalized to untreated proliferation control. Primer sequences are specified in the supplementary information (Table S1).

### FACS

Osteogenically differentiating hMSCs were trypsinized on day 21 and washed with Flow Cytometry Staining Buffer (FCSB, FC001, R&D systems) for two times. Cells were re-suspended in 100 μl of FCSB and 1 μl of mouse anti-human CD10-PE (561002, BD Biosciences) and mouse anti-human CD92-AF647 (565316, BD Biosciences) antibodies were added and incubated for 45 mins at 4 ⁰C. Cells were washed with FCSB twice and re-suspended in 200 μl of FCSB sorted by FACS (BD FACSAria II). FACS sorted cells were seeded on a glass coverslip in the osteogenic differentiation medium. Cells were transfected with SOX9-mGFP plasmid on day 22 and FRAP was performed on the 23^rd^ day of osteogenic differentiation.

## Results

### Interpretation of TF-FRAP results

We refer our readers to Govindaraj et al for the detailed explanation of TF-FRAP method and interpretation of data [13, 16]. In short: A higher immobile fraction (IF) and/or a longer recovery half-time of A_2_ (residence time) are the indicators of increased SOX9 transcriptional activity. Currently, there is no minimum cut-off value that defines the transcriptional output, but the values are always compared to those in other experimental conditions.

### Cellular heterogeneity leads to diverse SOX9 mobility patterns

Transcriptional activity of SOX9 and its target gene expression are known to be dynamically regulated during the chondrogenic, osteogenic and adipogenic differentiation of hMSCs [9, 17]. SOX9 activity is known to increase during chondrogenesis and cartilage homeostasis [18] and down regulated during osteogenic and adipogenic differentiation. Since SOX9 binding to DNA precedes its transcriptional activity, we studied its DNA binding dynamics in proliferating and differentiating hMSCs. We expected SOX9 binding to DNA and recovery half-times will be increased during chondrogenic differentiation and decreased during osteogenic and adipogenic differentiation, as compared to the rates obtained in undifferentiated MSCs? Data obtained by averaging TF-FRAP values per time point did not yield any meaningful information, except that the averaged SOX9 mobility was increased during differentiation into any lineage (Figure S1). We observed diverse SOX9 mobility patterns in our TF-FRAP measurements. Since MSCs are known to be a heterogeneous population of cells [19], we hypothesized that there may be different clusters of cells. For this, we performed hierarchical cluster analysis.

Unsupervised hierarchical clustering showed the presence of at least four clusters with distinct mobility patterns in undifferentiated hMSCs (Figure 1A and Table S2) as determined by immobile fraction and recovery half-times of SOX9. Cluster A had high IF and intermediate recovery half-times as compared to other clusters. Cluster B had high IF and low recovery half-times. Cluster C had intermediate IF and low recovery half-times. Cluster D had lower values for all three variables.

**Figure 1:**
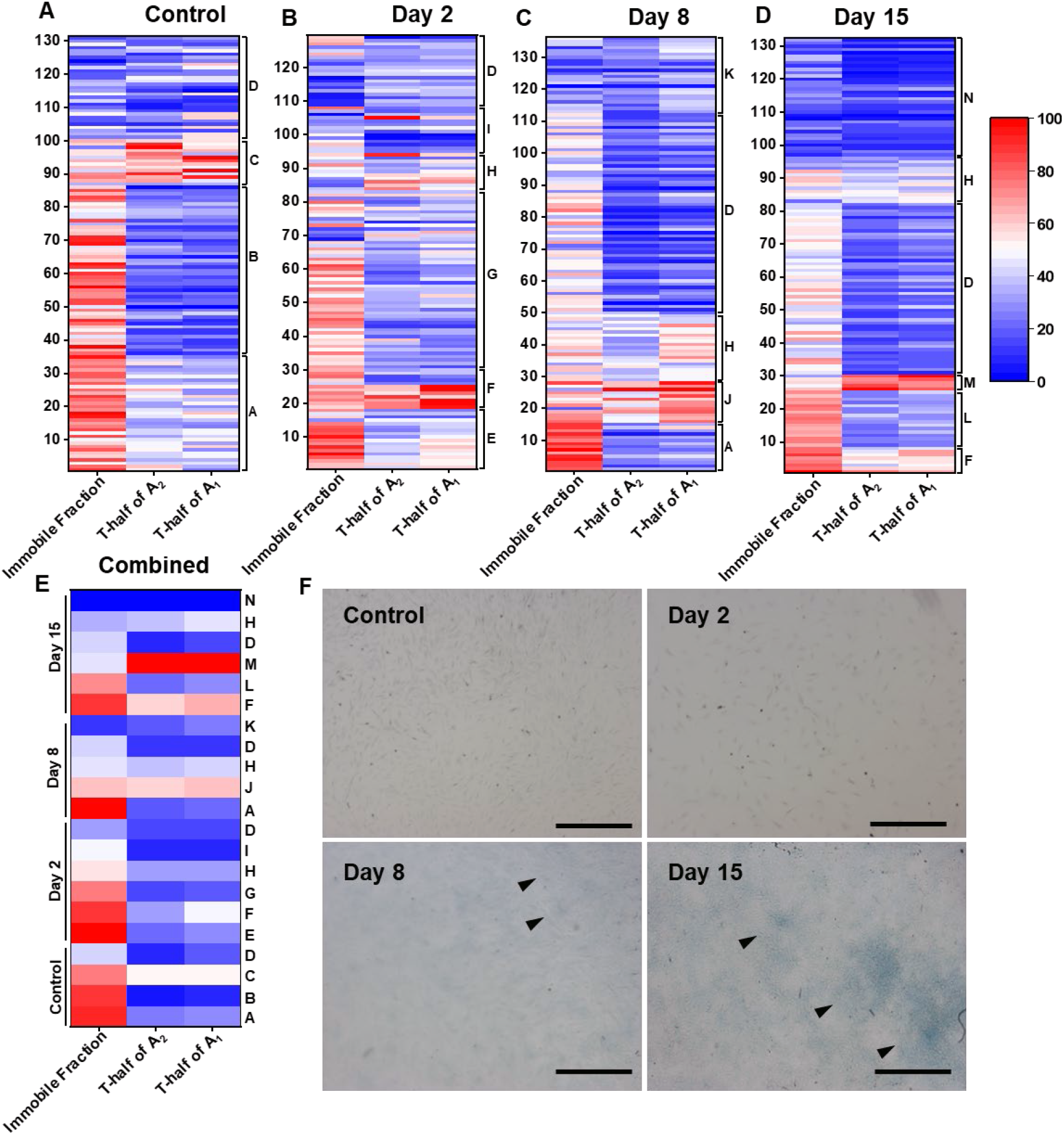
SOX9 mobility decreases during chondrogenic differentiation. **A-D**. Heat maps show the changes in SOX9 dynamics at the single cell level per cluster during the chondrogenic differentiation of hMSCs at day 0 (A), day 2 (B), 8 (C) and 15 (D). (E) Heat map showing averaged FRAP values per cluster across time-points, letters at the right indicate cluster ID. FRAP dynamics data from three hMSC donors were combined. FRAP dynamics data from three hMSC donors were combined. To fit in the heat map scale, all the variables were normalized (0, 100). N ≥ 129. (F) Alcian blue staining show an increase in GAG production during chondrogenic differentiation of hMSCs. Arrow heads indicate the Alcian blue stained GAG at day 2, 8 and 15. Scale bar: 500 μm.

TF-FRAP curves of individual clusters and relationship between IF and recovery half-time of A_2_ are presented in figure S2A-F. The percentage of cells per cluster is shown in figure S8A and table S2. To check whether these clusters in hMSCs are simply an integral property of any cell type, we measured the SOX9 dynamics in the C20/A4 cells and hPCs. TF-FRAP curves of SOX9 in the C20/A4 cells showed only one type of mobility pattern (Figure S3A), whereas hPCs showed two clusters with distinct mobility patterns (Figure S3B).

To rule out that the differential SOX9 dynamics patterns are due to various stages of the cell cycle, we performed FRAP measurements in undifferentiated hMSCs that were synchronized for 24 hours. Hierarchical clustering showed presence of at least four clusters in synchronized hMSCs as well (Figure S4). However, the number of cells per cluster was very different from those observed in unsynchronized hMSCs. This could be due to cell death during serum starvation and the number of cells that died per cluster could be different, depending on their metabolic state.

Although the population-doubling time for hMSCs is known to be about 30 – 39h [20], we could synchronize only for 24h as transfected cells started to die after that. In addition, we observed differential SOX9 mobility patterns in chondrogenically differentiating hMSCs, where cell division is limited due to lack of nutrients. Moreover, the well-established fact that hMSCs is a heterogenic population of cells, indicate that the diverse SOX9 mobility patterns in hMSCs are due to its heterogenic cell population. Thus, we applied unsupervised hierarchical clustering to cluster cells in all time-points and differentiation lineages.

### SOX9 residence time on DNA increases during chondrogenic differentiation

SOX9 activity is known to increase during chondrogenic differentiation of hMSCs as indicated by increased target gene expression [6]. To describe changes in SOX9 transcription dynamics, we measured its mobility by TF-FRAP during chondrogenic differentiation of hMSCs. SOX9 dynamics were measured at an early time point (day 2), at a middle time point (day 8) and at a later time point (day 15) and its mobility at those time points was compared with that of the undifferentiated hMSCs. After 15 days of chondrogenic differentiation cells were hard to transfect due to the presence of dense extracellular matrix around the cells. As this posed a possible transfection bias, by which some cells are transfected with higher efficiency than others, we used day 15 as the latest time-point.

During chondrogenic differentiation, we observed a total of 10 clusters (E-N) that were significantly different within. At day 2 and 15 of chondrogenic differentiation, we observed six clusters (Figure 1B and D) and five clusters at day 8 (Figure 1C) with distinct SOX9 dynamics. Interestingly, cluster D from the control group was present in all the three time-points of chondrogenic differentiation, indicating that at least one subpopulation of cells in hMSCs did not respond to the chondrogenic differentiation stimuli. Cluster H was present in all three differentiation time-points. Please note that the similarity of cluster D or H across time-points is not visible in the heat-maps, as the FRAP values were influenced in the normalization process by the appearance of new clusters. FRAP recovery curves of individual clusters and their relationship between IF and recovery half-time of A_2_ are presented in Figure S5A-F.

We expected increased SOX9 immobile fraction and longer recovery half times during chondrogenic differentiation as compared to undifferentiated hMSCs. We observed higher immobile fraction at day 2 and longer recovery half-times during day 8 and 15. Many of the new clusters that appeared in chondrogenically differentiating hMSCs showed a longer t-half of A_2_ (>12 sec) at day 15. Cluster M showed a t-half of A_2_ of more than 29 sec (Table S3). Our data show that during chondrogenic differentiation, the SOX9 dynamics changes differentially among the subpopulation of hMSCs. Comparing SOX9 dynamics at individual cluster level revealed that the SOX9-DNA binding (IF) decreased in most clusters during chondrogenic differentiation as compared to undifferentiated hMSCs (Figure 1E and Table S3). . Increased recovery half-times of SOX9 indicate longer residence times and active exchange of SOX9 on DNA. This is an indication that more chondrogenic genes under SOX9 control are differentially regulated during chondrogenic differentiation.

We confirmed the chondrogenic differentiation of hMSCs by Alcian blue staining (Figure 1F). The percentage of staining increased and visually correlated to the increase in differentiation at the later time-points. Increased immobility/lower diffusion times SOX9-DNA binding also correlated to increased COL2A gene expression (Figure S9A). The percentage of cells present per cluster during chondrogenic differentiation is presented in figure S8B.

### SOX9 binding and its residence time on DNA decreased during osteogenic differentiation

SOX9 activity is expected to be downregulated during osteogenic differentiation. Loebel *et al.* (2015) reported that SOX9 activity decreased (by means of decreased mRNA expression) during early stages (day 2 and 7) and is gradually increased at later stages (day 14 and 21) of osteogenic differentiation [21]. To further understand SOX9 protein activity during osteogenic differentiation, we studied SOX9 mobility during at day 2, 8 and 23 in osteogenically differentiating hMSCs. We observed 12 unique clusters (E – L) of SOX9 mobility patterns during osteogenic differentiation (Figure 2A-D).

**Figure 2:**
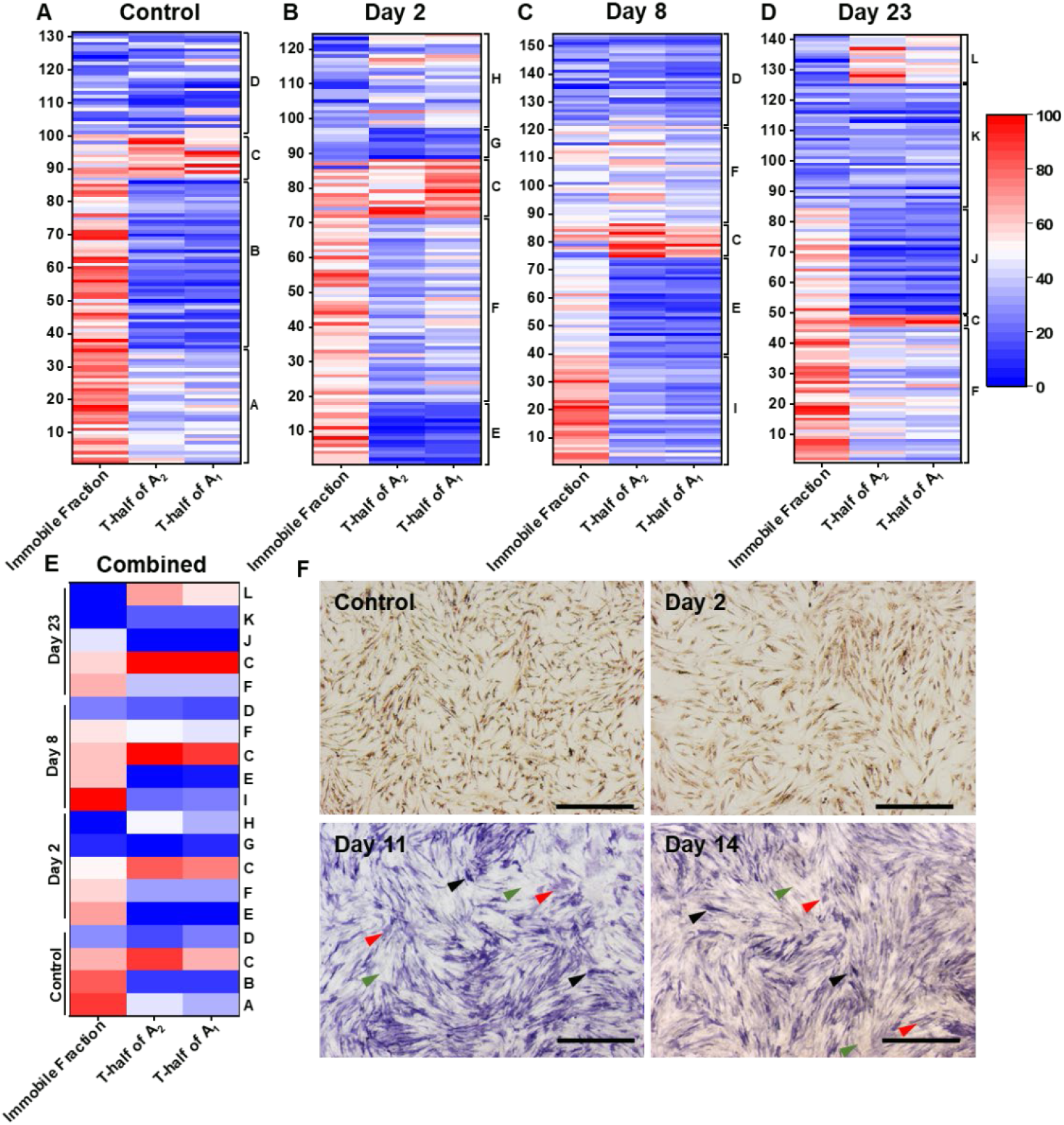
SOX9 binding decreases during osteogenic differentiation of hMSCs. (A-D) Heat map of FRAP rates show changes in SOX9 dynamics in undifferentiated hMSCs (A), at day 2 (B), at day 8 (C), and at day 23 (D). (E) Heat map showing averaged FRAP values per cluster across time-points, letters at the right indicate cluster ID. FRAP dynamics data from three hMSC donors were combined. To fit in the heat map scale, all the variables were normalized (0, 100). N ≥ 124. (F) ALP staining show an increase in ALP production during osteogenic differentiation of hMSCs. Black arrow heads indicate the cells with higher ALP production. Red arrow heads indicate the cells with lower ALP production. Green arrow heads indicate the cells with no ALP production. Scale bar: 500 μm.

We expected lower SOX9-DNA binding and/or recovery half-times during osteogenic differentiation as compared to undifferentiated hMSCs. Comparing SOX9 dynamics data at the single cell level shows that the SOX9-DNA binding (IF) decreased in many clusters during osteogenic differentiation as compared to undifferentiated hMSCs (Figure 2E). SOX9 binding was higher at day 2 as compared to day 8 and 23. However, the IF was relatively lower (except for cluster I, day 8) in differentiating hMSCs as compared to the undifferentiated hMSCs (Figure 2E).

Recovery half-times increased at day 8 and 23 as compared to day 2 of osteogenic differentiation. Interestingly, during osteogenic differentiation, recovery half-times of SOX9 were shorter than what is observed in the various clusters in chondrogenic differentiation. These data suggest that during osteogenic differentiation, SOX9 activity may be lower as compared to its activity during chondrogenic differentiation. FRAP curves of individual clusters of osteogenic differentiation at different time-points and their relationship between IF and recovery half-time of A_2_ are presented in the figure S6A-F.

Interestingly, cluster C from the undifferentiated hMSCs was present at all three differentiated time-points. However, the number of cells present in this cluster decreased during differentiation. Moreover, during differentiation, a new cluster F was present at all three time-points. In this cluster, IF was ∼30% and recovery half-time of A_2_ was > 10 sec. This cluster F contained considerably large amounts of cells (≥ 23%, Figure S8C). This suggests that cells in the cluster C may not differentiate into adipocytes and cells in cluster F may contain osteogenically differentiated cells.

ALP activity is one of the markers of osteogenic differentiation [22]. We performed ALP staining to confirm the osteogenic differentiation of hMSCs. As expected, ALP activity was higher at day 11 and 14. We did not observe ALP staining in the control group or at day 2. Notably, at day 11 and 14 we observed heterogenic ALP production in osteogenically differentiating hMSCs. Some cells produced high levels of ALP, while other cells showed lower levels and some cells had no ALP production (Figure 2F). This indicates that although all cells were exposed to the same differentiation stimuli, not all cells respond to the same extend. It suggests that there is a population of cells that did not undergo osteogenic differentiation, and those cells did not produce ALP (for example, clusters C and D). This partially explains our FRAP data in which SOX9 dynamics did not change in some clusters of differentiating hMSCs as compared to the control group (undifferentiated). The increased RUNX2 (at day 23) and *ALPL* gene expression correlated to osteogenic differentiation and SOX9-DNA binding (Figure S9B and C). All the FRAP rates for day 2, 8 and 23 of osteogenic differentiation are presented in tables S4.

### No changes in SOX9 dynamics patterns during adipogenic differentiation

SOX9 plays a minimal role in adipogenic differentiation of hMSCs [23]. We studied SOX9 dynamics in adipogenically differentiating hMSCs at day 2, 8 and 23. We expected the SOX9 immobile fraction and recovery half-times to be lower during adipogenic differentiation as compared to undifferentiated hMSCs. We observed a total of 9 (E-M) new clusters during adipogenic differentiation, four clusters at day 2, five clusters at day 8 and six clusters at day 23 (Figure 3A-D). Moreover, higher immobile fractions and longer recovery half-times were observed during adipogenic differentiation as compared to clusters in undifferentiated hMSCs (Figure 3A-D). There was also little change in SOX9 dynamics during adipogenic differentiation (Figure 3E) as compared to changes in other two differentiation lineages.

**Figure 3:**
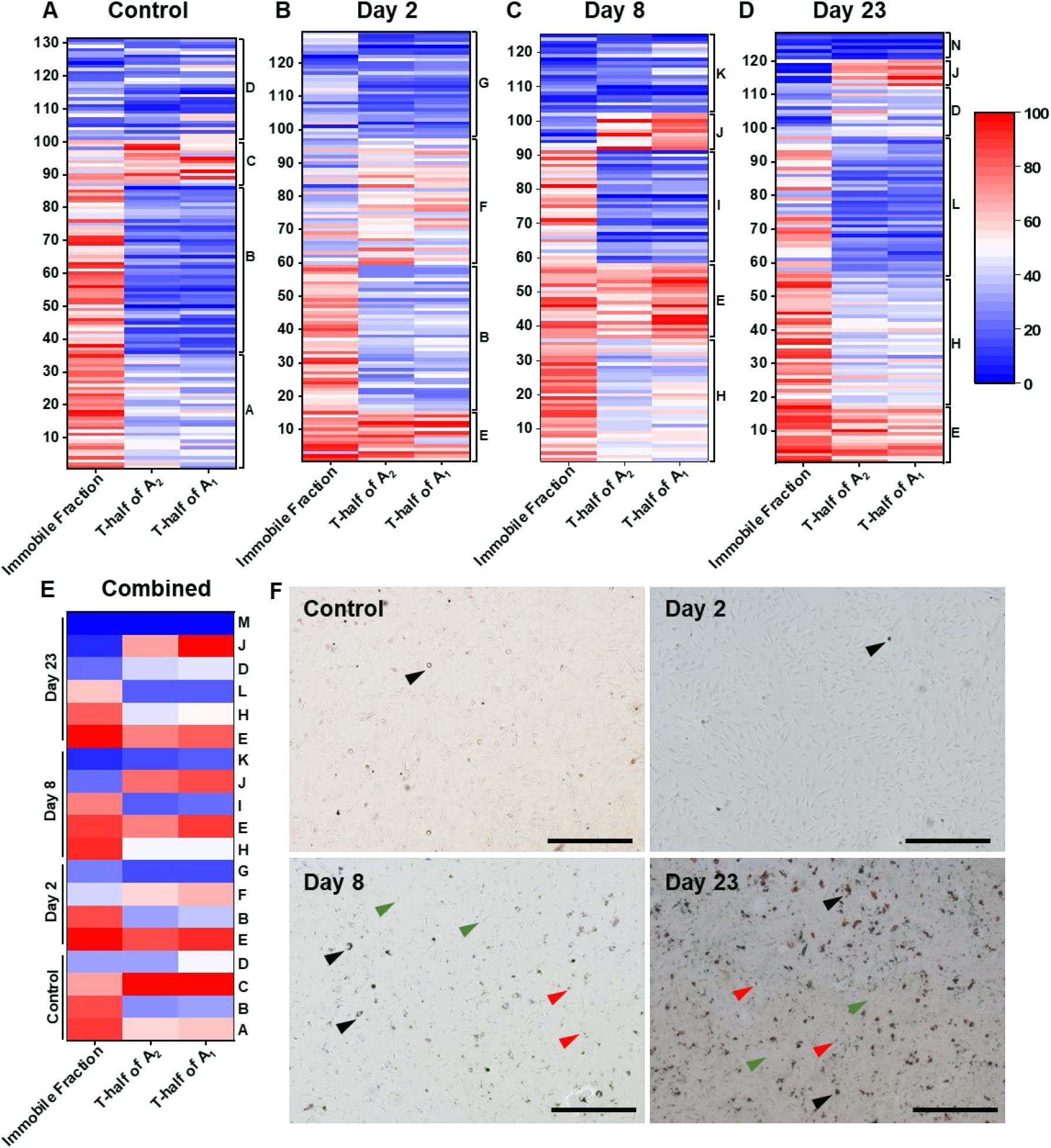
SOX9 residence time and binding decrease during the adipogenic differentiation of hMSCs. SOX9 dynamics in the undifferentiated hMSCs (A), day 2 (B), 8 (C) and 23 (D). (E) Heat map showing averaged FRAP values per cluster across time-points, letters at the right indicate cluster ID. FRAP dynamics data from three hMSC donors were combined. To fit in the heat map scale, all the variables were normalized (0, 100). N ≥ 125. (F) Oil Red O staining show an increase in lipid production during adipogenic differentiation of hMSCs. Black arrow heads indicate the cells with higher lipid production. Red arrow heads indicate the cells with lower lipid production. Green arrow heads indicate the cells with no lipid production. Scale bar: 500 μm.

A new cluster, named E, was present at all three differentiation time-points, although was found in only ∼14% of the cells. Cluster H was present at days 8 and 23, and it closely resembled cluster B, with very small changes in the FRAP variables. However, two FRAP variables in cluster B and H were significantly different and we therefore considered them to be separate clusters. The ratio of cells present in each cluster are shown in figure S8D and table S5.

Single cell SOX9 dynamics data show that SOX9 binding decreased in most cells. The dynamics pattern was quite similar across time-points (Figure 3A-D). FRAP recovery curves of individual clusters of adipogenic differentiation and their relationship between IF and recovery half-time of A_2_ are presented in the figure S7A-F. These data suggest that after initial changes, SOX9 mobility does not change dynamically within the clusters during the process of adipogenic differentiation.

We performed Oil Red O staining to stain the lipid droplets to confirm the adipogenic differentiation of hMSCs. We observed lipid droplets in a small population of undifferentiated hMSCs. However, the number of cells producing lipid droplets increased during adipogenic differentiation. At day 8 and 23 of adipogenic differentiation, we observed three types of cells, some had very intense staining with oil red O, some cells showed less staining for lipid droplets and a third group had no visible lipid droplets (Figure 3F). This suggests that although all cells were exposed to the same differentiation stimuli, not all cells differentiated into adipocytes. qPCR analysis revealed elevated PPARγ gene expression, which correlates to the adipogenic differentiation of hMSCs (Figure S9D).

### SOX9 residence time on DNA influence its target gene expression

We compared SOX9 residence time and DNA binding between chondro-, osteo- and adipogenic differentiation, at the subpopulation level. We averaged TF-FRAP rates within the subpopulation and normalized (0, 100) across time-points and differentiation. We expected SOX9 residence times and DNA binding would be higher in clusters of chondrogenically-differentiated hMSCs as compared to other two differentiation lineages. However, we observed the expected changes only in the SOX9 residence time. SOX9 residence times were the highest in the clusters of chondro-, modest in the clusters of osteo- and the lowest in the clusters of adipogenic differentiation. Although, we observed clusters with higher immobile fraction in the chondrogenically-differentiated hMSCs, some clusters in adipogenically-differentiated hMSCs also contained comparatively higher immobile fractions. These data suggest that SOX9 transcriptional activity may be comparatively higher during chondro-, moderate during osteo- and lower during adipogenic differentiation (Figure 4A). Interestingly, changes in SOX9 residence time on DNA corrlated to its target gene expression levels. SOX9 gene expression itself was higher during chondorgenic differentiation (at day 8 and 15) as compared to other differentiation lineages. COL2A (target gene of SOX9) expression was higher at day 8 and 15 of chondrogenic differentiation, where, SOX9 had longer residence time. With comparatively longer residence time of SOX9, COL2A expression was also higher during osteogenic differentiation as compared to adipogenic differentiation (Figure 4B). This indicates that the longer residence time of SOX9 on DNA results in higher transcriptional output.

**Figure 4:**
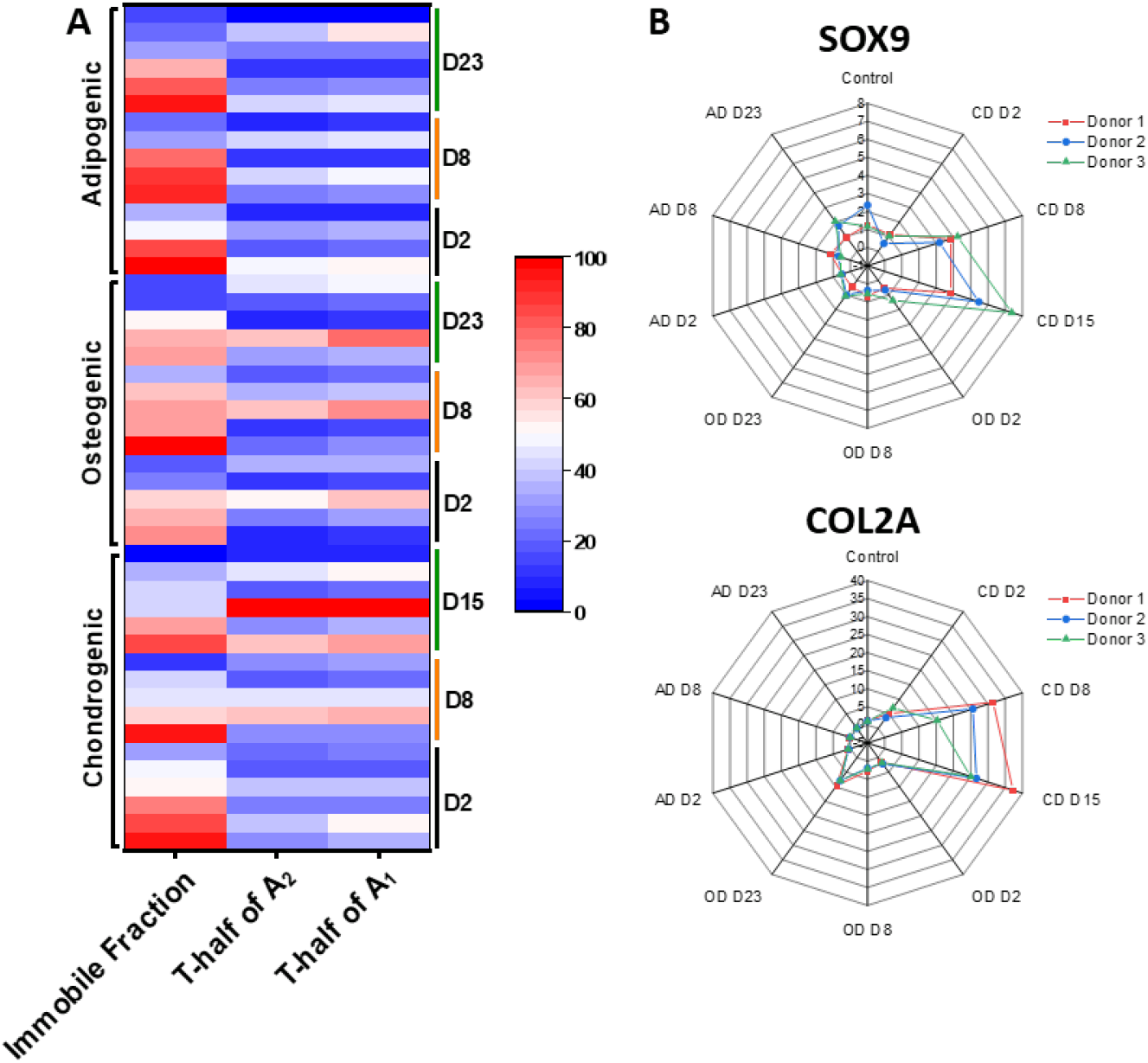
SOX9 residence time on DNA decreased from chondrogenic to osteogenic to adipogenic differentiation, which correlate to gene expression. (A) Heat map comparing FRAP rates of SOX9-mGFP during chondro-, osteo-, and adipogenic differentiation, averaged at subpopulation level. Clusters in chondrogenic differentiation show longer Recovery half-times as compared to other differentiation lineages. Clusters in adipogenic differentiation show shorter recovery half-times as compared to other two differentiation lineages. Clusters in osteogenic differentiation show shorter recovery half-times as compared to chondrogenic and longer as compared to adipogenic differentiation lineages. Immobile fraction was higher in the initial stages chondrogenic differentiation and later stages on adipogenic differentiation. (B) Gene expression analysis by qPCR show that SOX9 and its target gene COL2A expression is increased at day 8 and 15 of chondrogenic differentiation as compared to other differentiation and time-points. During osteogenic differentiation, COL2A expression was higher than adipogenic differentiation but lower than chondrogenic differentiation (qPCR: n = 3 donors, with triplicates in each donor)

### Differential dynamics of SOX9 are the result of hMSC heterogeneity

Osteogenically differentiated hMSCs are known to be positive for CD10 and CD92 cell surface markers [24]. To confirm that the heterogenic SOX9 dynamics in hMSCs is due to cellular heterogeneity, we sorted osteogenically differentiated hMSCs (at day 23) for surface markers positive for either CD10 or CD92 or both, and performed FRAP on these cells (Figure 5A-E). We expected FRAP rates of FACS sorted cells for CD10+ and/or CD92+ would be homogeneous as they are osteogenically differentiated single sub-population of cells. As expected, these FACS sorted cells showed less heterogeneity in cells that are positive for either CD10+ or CD92+ (Figure 5A and B) and a nearly homogeneous SOX9 mobility pattern is observed in double positive (CD10+ and CD92+) cells (Figure 5C). Recovery half-times of A_1_ and A_2_ of osteogenically differentiated CD10+ cells were not significantly different from cluster K of osteogenically differentiated hMSCs at day 23. The FRAP values are shown in the table S16.

**Figure 5:**
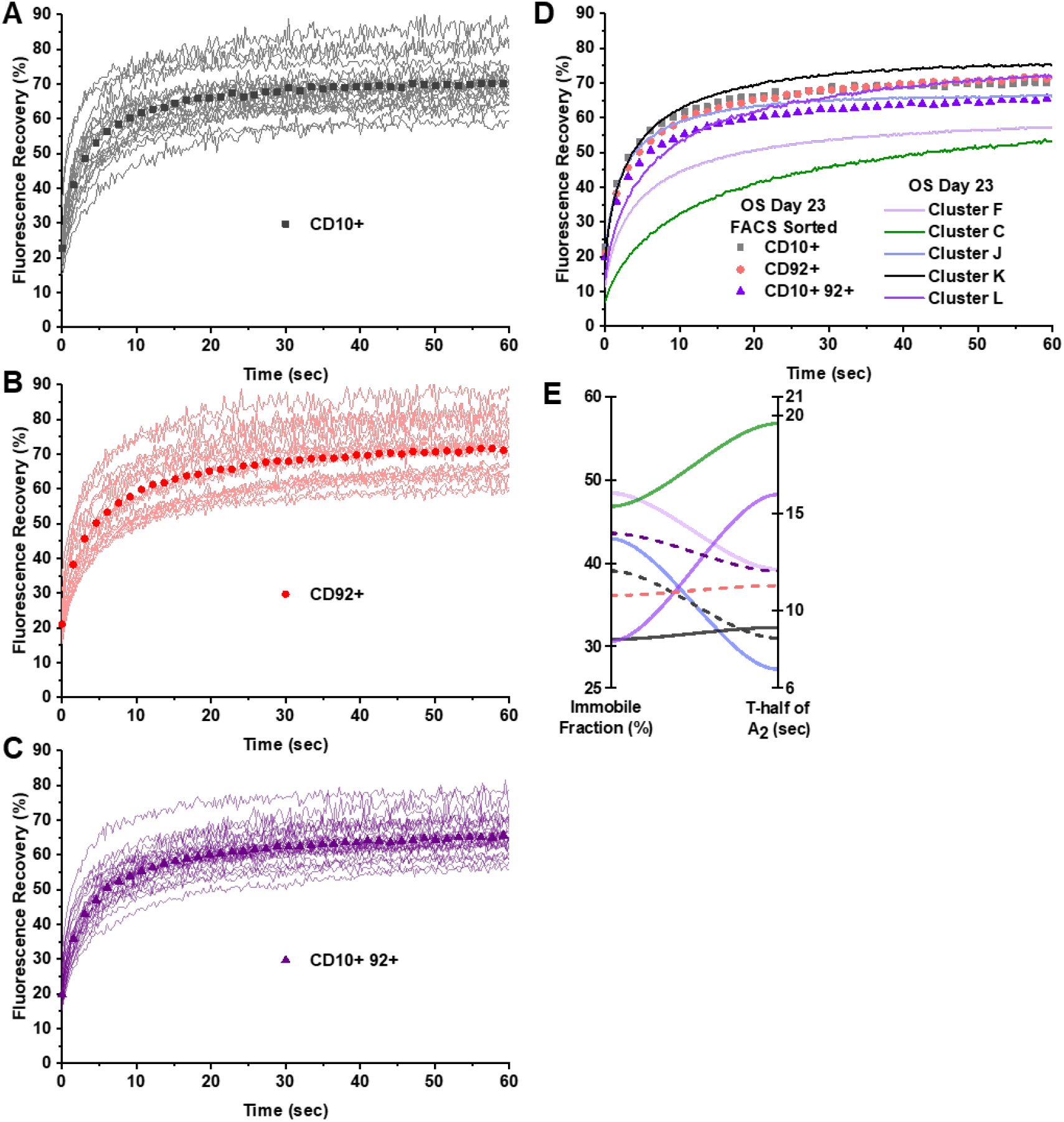
FACS sorted osteogenically-differentiated hMSCs (day 23) show differential mobility of SOX9 is due to hMSC heterogeneity. Osteogenically differentiated hMSCs were sorted for CD10+ or CD92+ or both markers and SOX9 mobility was measured by FRAP. SOX9 mobility in FACS sorted cells show less heterogeneity. SOX9 mobility in (A) CD10+, (B) CD92+ and (D) CD10+ and CD92+ cells of osteogenically differentiated hMSCs. FRAP curves of continuous line show recovery in individual cells and symbols (square, circle and triangle) show the average. (F) Averaged FRAP curves of CD10+ and/or CD92+ are compared to osteogenically differentiated hMSCs (day 23). (E) Relationship between immobile fraction and t-half of A_2_ is compared between FACS sorted (dashed lines) and Unsorted (continuous lines) cells. n ≥ 19 per condition.

Absence of heterogeneity in double positive cells (CD10 and CD92) indicates that a subpopulation of cells in hMSCs have distinct SOX9 dynamics because of the cellular heterogeneity. Differences in FRAP rates between CD10+ and CD92+ cells were very small as compared to CD10+ / CD92+ double positive cells. We performed this experiment in only one donor and compared with the relevant cluster of osteogenically differentiated cells (day 23). FACS sorted and unsorted cells were compared at day 23 of osteogenic differentiation. Therefore, the differences in the FRAP rates (between FACS sorted and unsorted cells) may be because FACS sorted cells were subjected to more stress, such as antibody staining and mechanical stress during FACS sorting. We chose osteogenically differentiated hMSCs over other lineages due to its high transfection efficiency.

### SOX9 dynamics in chondrogenically-differentiated hMSCs did not resemble healthy or OA hPCs

SOX9 drives chondrogenic differentiation and maintains cartilage homeostasis. Healthy articular chondrocytes are resistant to hypertrophic differentiation. However, during the onset of osteoarthritis, these healthy articular chondrocytes undergo hypertrophic differentiation. During this time SOX9 activity is known to decrease and RUNX2 activity increases [3, 25]. However, this hypertrophic switch is not yet fully understood. To gain more insight into SOX9 dynamics during differentiation, we compared the SOX9 dynamics of healthy and OA hPCs with mobility patterns of undifferentiated and chondrogenically-differentiated hMSCs (day 15). We expected to find chondro progenitor cells in undifferentiated hMSCs by comparing its SOX9 dynamics with those of healthy hPCs. In hPCs we identified two clusters of cells, as described before (unpublished). Two clusters in the undifferentiated hMSCs (cluster A and C) visually closely resembled cluster 2 of healthy hPCs (Figure 6A, B). Recovery half-times of A_1_ and A_2_ were not significantly different between cluster A of undifferentiated hMSCs and cluster 2 of healthy hPCs. Although the difference in the IF between cluster A and cluster 2 was less than 5%, the change was statistically significant (Table S7). This indicates that the cells in cluster A of undifferentiated hMSCs – resembles the cluster 2 of healthy hPCs – might have a higher chondrogenic potential as compared to cells in other clusters.

**Figure 6:**
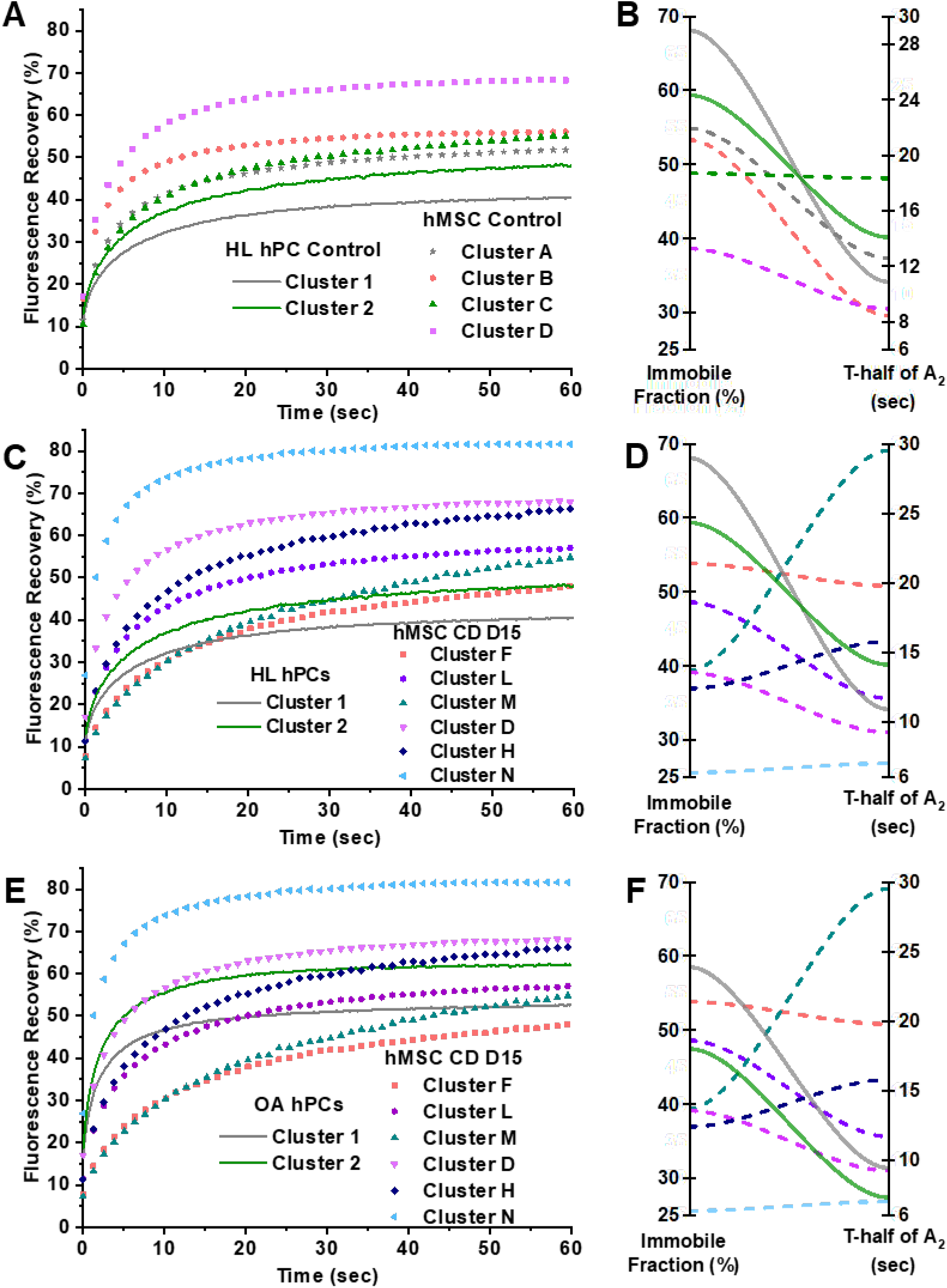
SOX9 mobility patterns in 15-day chondrogenically-differentiated hMSCs resemble neither healthy nor OA chondrocytes. SOX9 mobility is comparted between healthy hPCs and undifferentiated hMSCs (A and B). SOX9 mobility is comparted between healthy hPCs and chondrogenically-differentiated hMSCs at day 15 (C and D). SOX9 mobility is comparted between OA hPCs and chondrogenically-differentiated hMSCs at day 15 (C and D). n ≥ 40 per condition. hMSC control: undifferentiated hMSCs, CD: chondrogenic differentiation, D15: Day 15.

We also compared SOX9 dynamics of chondrogenically differentiated hMSCs (day 15) with healthy and OA hPCs. We expected to find chondrogenically differentiated clusters by comparing the SOX9 dynamic rates with those of healthy hPCs or OA hPCs, in case of hypertrophic differentiation. However, none of the clusters in the chondrogenically-differentiated hMSCs (day 15) resembled the SOX9 clusters of either healthy or OA hPCs (Figure 6C-F). This suggests that after 15 days of chondrogenic differentiation of hMSCs in monolayer no mature healthy chondrocytes are formed. This is also visible by the relatively low Alcian Blue staining as well as by the low COL2A1 expression at this time-point.

### Nuclear localization pattern of SOX9 is distinct between clusters

We observed distinct nuclear localization patterns of SOX9 in the control (undifferentiated) and in differentiating hMSCs. We checked whether these localization patterns were consistent within a cluster. Over 75% of cells in a cluster had distinct nuclear localization patterns unique for that cluster, as presented in figure 7. Cluster A showed a lower mobility of SOX9 as compared to other clusters in the control, and this cluster had comparatively more a discrete SOX9 localization. In cluster B and C we observed more punctate foci, especially in the nuclear periphery. Nuclei in cluster D showed a more homogeneous distribution of SOX9 and showed higher SOX9 mobility. A montage of nuclei showing SOX9 localization patterns of clusters A-D is presented in supplemental figures S10-S13.

**Figure 7:**
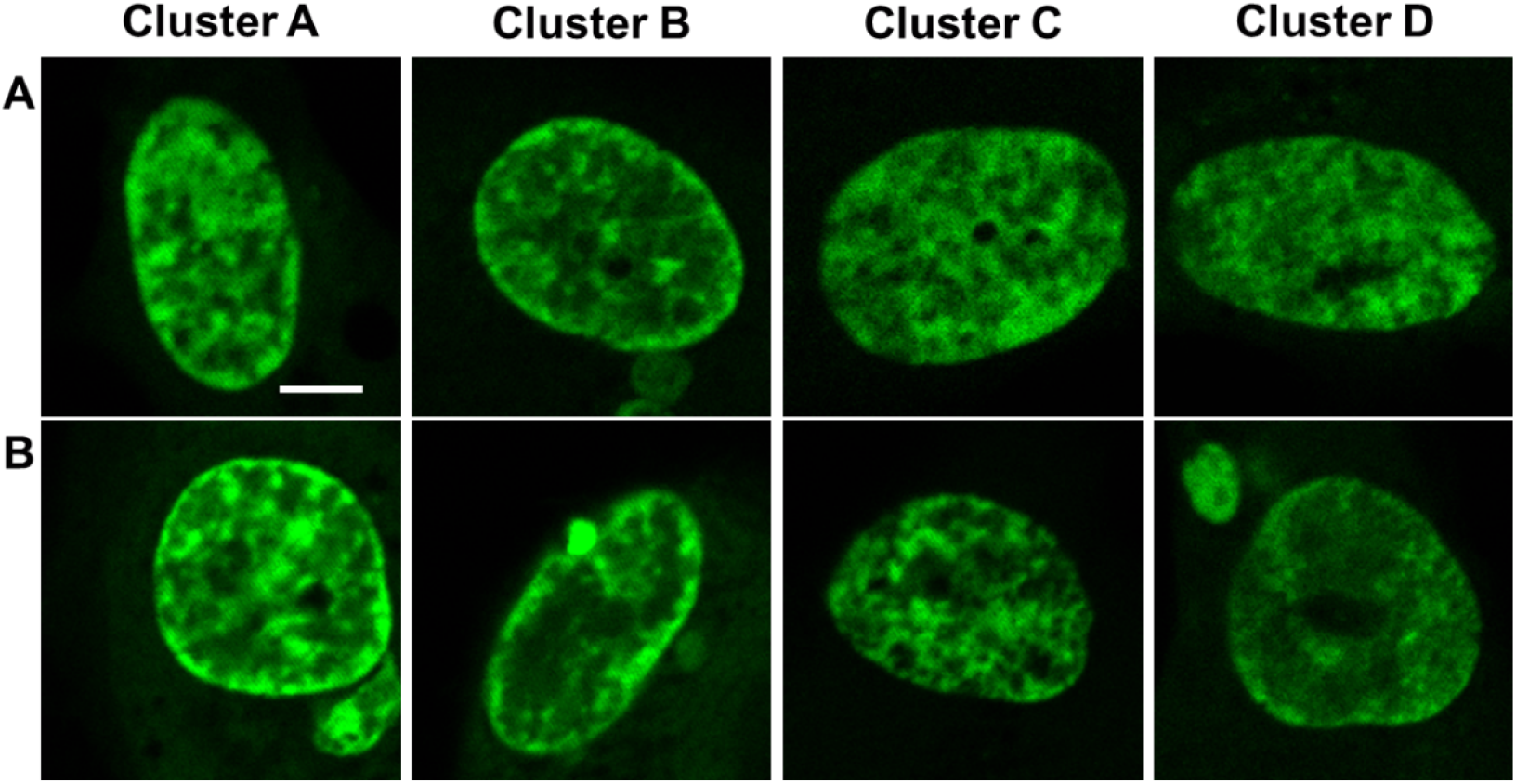
SOX9-mGFP show distinct nuclear localization patterns in subpopulations of hMSCs. Two representative cells from each cluster of undifferentiated hMSCs are shown (A and B). Cells in cluster A exhibit discrete SOX9-mGFP localization patterns and comparatively higher expression. Cells in cluster B and C exhibit either patched or homogeneous SOX9-mGFP localization patterns. Cells in cluster D exhibit diffused SOX9-mGFP localization pattern, which results in a higher mobility with lower IF and shorter t½ of A_2_. Scale bar: 5 μm.

To determine if various stages of the cell cycle contributed to the differential nuclear localization patterns of SOX9, we performed a cell cycle synchronization study by serum starving the cells for 24 hours, as described before [26]. We transfected the cells with SOX9-mGFP plasmid and supplemented with media containing FBS for about 8 - 10 hours to allow the cells to express SOX9-mGFP protein. We then starved the cells by replacing the media without FBS for 24 hours and observed the cells in the microscope. After 24 hours, we again replaced with media with FBS for 6 hours and imaged the cells at 30 hours (24h + 6h). After 30 hours of starting synchronization, too many transfected cells started dying, so we could not image at later time-points. After cell cycle synchronization, similar SOX9 nuclear localization patterns were observed, making the possibility of cell cycle playing a role in this less likely (Figure S14).

Although cell cycle synchronization for 24h is short as compared to cell doubling time of hMSCs (∼30-40h), we have observed differential SOX9 pattern at all time points of chondrogenically differentiating hMSCs as well, which are synchronized by addition of Dexamethasone, and wherein cell division is limited. To confirm that these patterns were not due to artifacts of overexpression of SOX9-mGFP, we performed immunofluorescence assays for endogenous SOX9 in untransfected hMSCs. Immunostaining confirmed that these nuclear localization patterns were inherent to the hMSCs (Figure S15). These data indicate that the spatial arrangement of SOX9 is different in subpopulations of hMSCs, and that this correlates to its mobility. Since SOX9 binding to DNA is linked to its transcriptional activity, the differential nuclear localization patterns suggest a differential gene expression in these clusters.

### The number of progenitor cells might determine differentiation potential of the donor

We assessed whether the differentiation potential of a donor towards a particular lineage (chondrogenic or osteogenic or adipogenic lineage) can be determined based on the number of progenitor cells present per cluster within the donor. For this, we assessed the differentiation potential of three donors towards a particular lineage based on the histology staining (GAG, ALP and Oil Red O staining) and compared it with the number of progenitor cells present per cluster within the donor (Figure 8).

**Figure 8:**
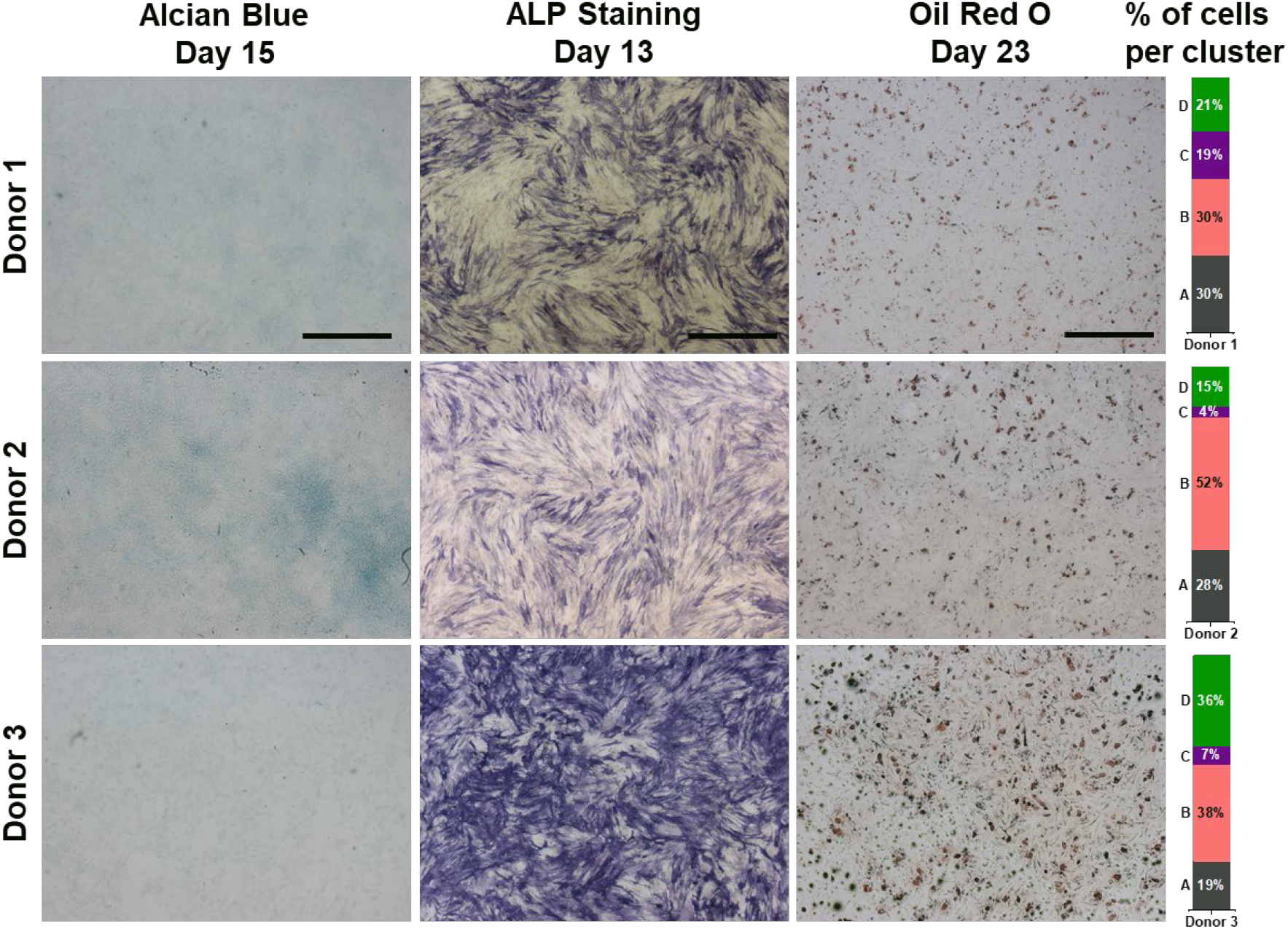
Number of progenitor cells present per cluster in the undifferentiated hMSCs determine the differentiation potential of the donor. Stacked column chart at the right indicate the percentage of cells per cluster and the cluster ID is indicated at the left of the column chart. Donor 1 had considerable amount of cells in each cluster and showed moderate differentiation potential towards all lineage as compared to other donors. Donor 2 had more chondrogenic potential as compared to other two donors as evidenced by more GAG staining. This donor had 52% of cells in the cluster B, which had higher IF of SOX9. Donor 3 had more osteogenic and adipogenic potential as compared to other two donors as evidenced by more ALP and Oil Red O staining respectively. This donor had higher number of cells in the cluster B (38%) and D (36%), the later had lowest IF and shorter t-half of A_2_ of SOX9. Scale bar: 500 μm.

Combined (cells from three donors) cluster analysis showed the presence of at least four clusters in the undifferentiated hMSCs. However, every donor had varying numbers of cells per cluster. Currently, based on the SOX9 dynamics, it is not possible to assign a particular cluster to be the progenitor cells for a particular lineage differentiation. We postulate that the presence of a higher number of cells in a particular cluster within a donor might result in a higher differentiation potential towards a particular lineage.

Interestingly, donor 1, which had a relatively even distribution of cells per cluster (20-30%), showed a moderate differentiation potential towards all lineages. Donor 2, which had a higher number of cells (52%) in cluster B, had a higher chondrogenic potential than the other two donors (1 and 3). Cluster A contained the cells with lower SOX9 mobility, i.e., higher IF and longer t-half of A_2_. The lower mobility of SOX9 indicates a higher binding of SOX9 to DNA, which results in more target (chondrogenic) gene expression [13]. In contrast, donor 3 had a higher number of cells in two clusters B (38%) and D (36%), and showed a higher osteogenic and adipogenic potential as compared to the other two donors (1 and 2). Cluster D contained the cells with higher SOX9 mobility, i.e., a lower IF and a shorter t-half of A_2_. The higher mobility of SOX9 indicates a lower binding to DNA resulting in lower target gene expression. These data suggest that the number of cells per cluster may be indicative of the presence of different progenitor cells and thus indicative of the differentiation potential of the donor. However, due to the large donor variation more donors will need to be tested to confirm this finding.

## Discussion

hMSCs are a promising cell type for regenerative medicine, due to its multi lineage differentiation potential. However, lack of sufficient knowledge of the signaling interplay and the role of cellular heterogeneity during differentiation, prevent the successful application of hMSCs in regenerative therapy [2]. We have previously shown that the SOX9-DNA binding directly linked to its transcriptional activity. Higher immobile fractions or longer recovery half-times resulted in an increase of expression of its target genes [13]. Here, we show that SOX9 dynamics and coupled transcriptional activity is different among subpopulations of hMSCs and that TF-FRAP can be used to determine the number of subpopulations in heterogenic cells and possibly the differentiation potential of a donor.

SOX9 gene and protein expression is known to increase during chondrogenic differentiation [6] and decreases during osteogenic differentiation of hMSCs [21]. There are several reports stating that hMSCs are comprised of heterogenic cell populations that differ in molecular phenotype is [27–29]. However, these differentiation studies were performed in hMSCs in bulk experiments, with little information at the subpopulation level. Moreover, gene and protein expression alone does not reveal their activity. To our knowledge, we are the first to show the dynamic changes of SOX9 protein activity at the subpopulation level in the differentiating hMSCs at the single cell level. Changes in SOX9 dynamics were high among the clusters during chondrogenic and osteogenic differentiation, as compared to adipogenic differentiation. This is in line with previous reports that show that SOX9 plays a key role during chondrogenic and osteogenic differentiation, but a lesser role during adipogenic differentiation [23].

Previous reports have shown that longer residence time (recovery half-time) of transcription factors result in higher target gene expression [30, 31], and we have shown this specifically for SOX9 [13]. During chondro- and osteogenic differentiation, many clusters showed higher residence time at day 8 and 15 as compared to day 2, indicating higher SOX9 transcriptional activity. However, overall residence time was higher during chondrogenic differentiation as compared to osteogenic differentiation. Which resulted in a higher COL2A expression during chondro- and osteogenic differentiation as compared to undifferentiated and adipogenically differentiating hMSCs. This is in line with existing studies based on RUNX2/SOX9 mRNA expression [21]. During adipogenic differentiation, although SOX9-DNA binding was higher, residence times were much lower than seen in the cells differentiating towards the other two differentiation lineages. This suggests that there is less active exchange of SOX9 at its binding sites, which would result in lower transcriptional output. Gulyaeva et al (2018) has shown that inactivation of Sox9 is a prerequisite for adipogenesis [32].

During chondrogenic, osteogenic and adipogenic differentiation of hMSCs, at least one cluster had a comparatively low immobile fraction and a shorter recovery half-time of A_2_ in at least 10% of cells. In contrast, there were also new clusters with either a higher immobile fraction or longer recovery half-times of A_2_, or both. Notably, in every differentiation lineage, at least one other cluster did not respond to the differentiation stimuli. For example, cluster D during chondrogenic differentiation, cluster C during osteogenic differentiation and cluster B (at day 2) during adipogenic differentiation did not respond to differentiation stimuli. It is interesting to note that different subpopulations did not respond to a particular differentiation lineage, indicating subpopulations in undifferentiated hMSCs have commitment towards a particular lineage.

Varying levels of ALP and Oil Red O staining at the single cell level during osteo- and adipogenic differentiation, indicate that individual cells have varying degree of differentiation potential. Some cells show higher, some show moderate, while others show lower-to-none ALP/lipid droplet production. Cells with higher ALP/lipid production indicate that they had a higher osteogenic/adipogenic potential, and vice versa. The observation that there are cells that showed no ALP/lipid production, may be indicative of the lower-to-none osteogenic/adipogenic potential.

Differential SOX9 dynamics as measured by TF-FRAP in hMSCs (undifferentiated/differentiated) can also be explained by differential ALP and lipid production at the single cell level. Individual clusters have varying differentiation potential towards a particular lineage, which is determined by the activity and coupled dynamics of transcription factors. In a clonal selection study, Prins et al (2014) has shown that individual clones (derived from single cells of hMSCs) from the same donor had varying differentiation potentials towards a particular lineage differentiation [33]. Selective clones showed tri lineage (chondro-, osteo- and adipogenic), two lineage, one lineage, or no lineage differentiation potential. In another study, Leyva Leyva et al (2012) isolated two subpopulations (CD105+ and CD105-) from the same hMSC donor and showed that these populations had varying levels of osteogenic differentiation potential [34]. Our group has previously shown that differentiation potential is coupled to specific miRNA expression patterns [35]. These studies strengthen our data that the subpopulations – as determined by SOX9 dynamics and ALP/lipid production at the single cell level – have varying differentiation potentials towards a particular lineage.

Cluster analysis of SOX9 dynamics identified at least four subpopulations (cell types) in the undifferentiated hMSCs and four to six subpopulations in the differentiating hMSCs, independent of the lineage of differentiation. Interestingly, the percentage of cells present per cluster in the undifferentiated hMSCs were different among the donors. These donors also had varying differentiation potential towards a particular lineage. Donor 1 and 2 had more than 60% cells in clusters A and B, and showed comparatively moderate and higher chondrogenic potential, respectively. Cluster D, which contains cells with lower SOX9-DNA binding and residence time, contained a higher number of cells in donor 3 as compared to other donors and showed higher osteogenic and adipogenic potential. This might reflect the influence of the number of cells in a progenitor subpopulation to the differentiation potential of a donor towards a particular lineage. Previously, Prins et at (2014) suggested predicting the differentiation potential of a donor based on ALP production at day 15 [33].However, our TF-FRAP might provide a novel technique to determine the differentiation potential of a donor towards a particular lineage.

The SOX9 dynamics of cells in cluster A of undifferentiated hMSCs were similar to those of cells in cluster 2 of healthy hPCs, indicating that this cluster might have a higher chondrogenic potential. Many reports have shown that the chondrogenic differentiation of hMSCs in 2D culture is less efficient as compared to 3D culture [36, 37]. Moreover, in vitro chondrogenic differentiation of hMSCs mimic hypertrophic chondrocytes, which are unsuitable for clinical applications [38]. In line with previous reports, our data suggest that SOX9 dynamics in chondrogenically differentiated hMSCs more closely resemble those of hypertrophic hPCs.

One of the limitations of our study, may be that there are differences in transfection efficiency between different cell types in the heterogeneous population of MSCs. It is known that transfection efficiency can be different among various cell types [39]. One should be aware that our findings on the percentage of cells that are present per cluster, based on the SOX9 dynamics, may be slightly variable if the transfection efficiency is not same among the clusters.

SOX9 target gene expression is known to be different among various cell types [13, 40, 41]. It is also interesting to note that the comparison of SOX9 dynamics between C20/A4, hPCs and subpopulation of hMSCs indicates that the dynamics of the same transcription factor is different among different cell types and in different subpopulations of heterogenic cells. This supports our previous finding that the activity of a transcription factor is dependent on its dynamics [13]. The number of cell types/subpopulations present in the hMSCs are unknown [29]. Tedious and complicated single cell RNA sequencing is one of the ways to identify subpopulations [42]. TF-FRAP identified at least four subpopulations in hMSCs with distinct SOX9 dynamics in the undifferentiated hMSCs. Our finding that cells expressing a particular or combination of CD markers present more homogeneous SOX9 dynamics, suggests that TF-FRAP can be used to identify the number of subpopulations present in heterogenic cells, such as hMSCs. Simple TF-FRAP on a master transcription factor and subsequent cluster analysis might reveal the number of subpopulations present in any heterogenic population.

The higher spatio-temporal resolution of TF-FRAP as a technique, has shed more insight into correlation between the spatial arrangement of SOX9 in the nucleus and its corresponding dynamics. The distinct nuclear localization patterns of SOX9 in these subpopulations of hMSCs correlated to its differential dynamics pattern. It was a serendipitous observation that a transcription factor can localize differentially in the nucleus in the different subpopulations of hMSCs, and that this resulted in differential mobility patterns. Cells with strong SOX9 expression and/or discrete foci showed comparatively higher DNA binding and longer residence time and *vice versa*. The presence of discrete subnuclear foci of a transcription factor is known to show higher transcriptional output of target genes [43–45]. It is tempting to say that the cells that were present in cluster A, that had higher SOX9 DNA binding, a higher residence time of SOX9 coupled to discrete subnuclear foci, and that are similar to one of the clusters in hPCs, might be the chondro-progenitor cells.

Together, we have shown that subpopulations of hMSCs have differential SOX9 dynamics. SOX9 transcriptional activity is different among these subpopulations as evidenced by varying residence time, GAG/ ALP/lipid droplet production and differential subnuclear localization patterns. TF-FRAP can prove to be a novel method to predict the number of subpopulations present in a heterogenic cells and predict the differentiation potential of a donor based on transcription factor dynamics.

### Author contributions

KG and SK designed and performed the experiments and analyzed the data. JP conceived the idea and designed the experiments. MK critically revised the manuscript. All authors contributed to the writing of the manuscript.

