## Supplementary material for "Mapping SOX9 transcriptional dynamics during multi-lineage differentiation of human mesenchymal stem cells": Figure S8

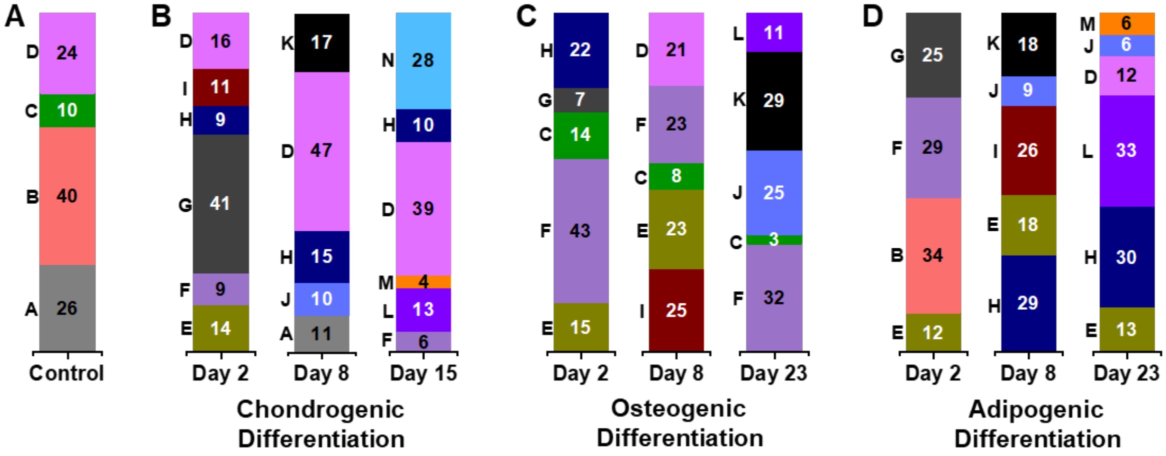


***Figure S8: Number of cells present in each cluster is variable depending on the differentiation lineage and time.*** *Bar graphs show proportion of cells in present in each cluster of the undifferentiated hMSCs (A), chondrogenically (B), osteogenically (C) and adipogenically (D) differentiating hMSCs. Letters to the left of the bar graph indicate the cluster ID. Number of cells present in the differentiated clusters increased in day 8 and 15 or 23 as comparted to day 2.*
