## Supplementary material for "Mapping SOX9 transcriptional dynamics during multi-lineage differentiation of human mesenchymal stem cells": Figure S9

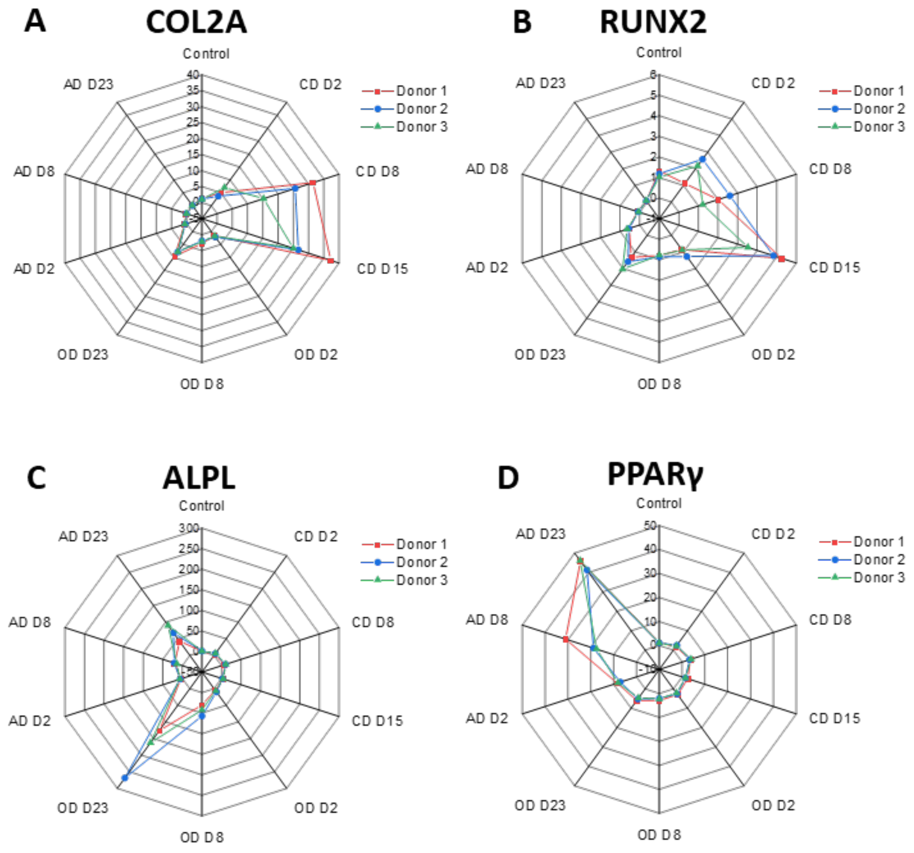


***Figure S9: Gene expression profile confirm the chondro, osteo and adipogenic differentiation of hMSCs.*** *Spider plots show mRNA expression for (A) COL2A, (B) RUNX2, (C) ALPL and (D) PPARγ for all differentiation lineage and time points. Data is shown for all three donors, qPCR was performed in triplicate per donor. Control is undifferentiated hMSCs at day 0. D = Day, CD = chondrogenic, OD = osteogenic and AD = adipogenic differentiation, Scale is fold change, normalized to RPL13.*
