## Supplementary material for "Mapping SOX9 transcriptional dynamics during multi-lineage differentiation of human mesenchymal stem cells": Figure S10

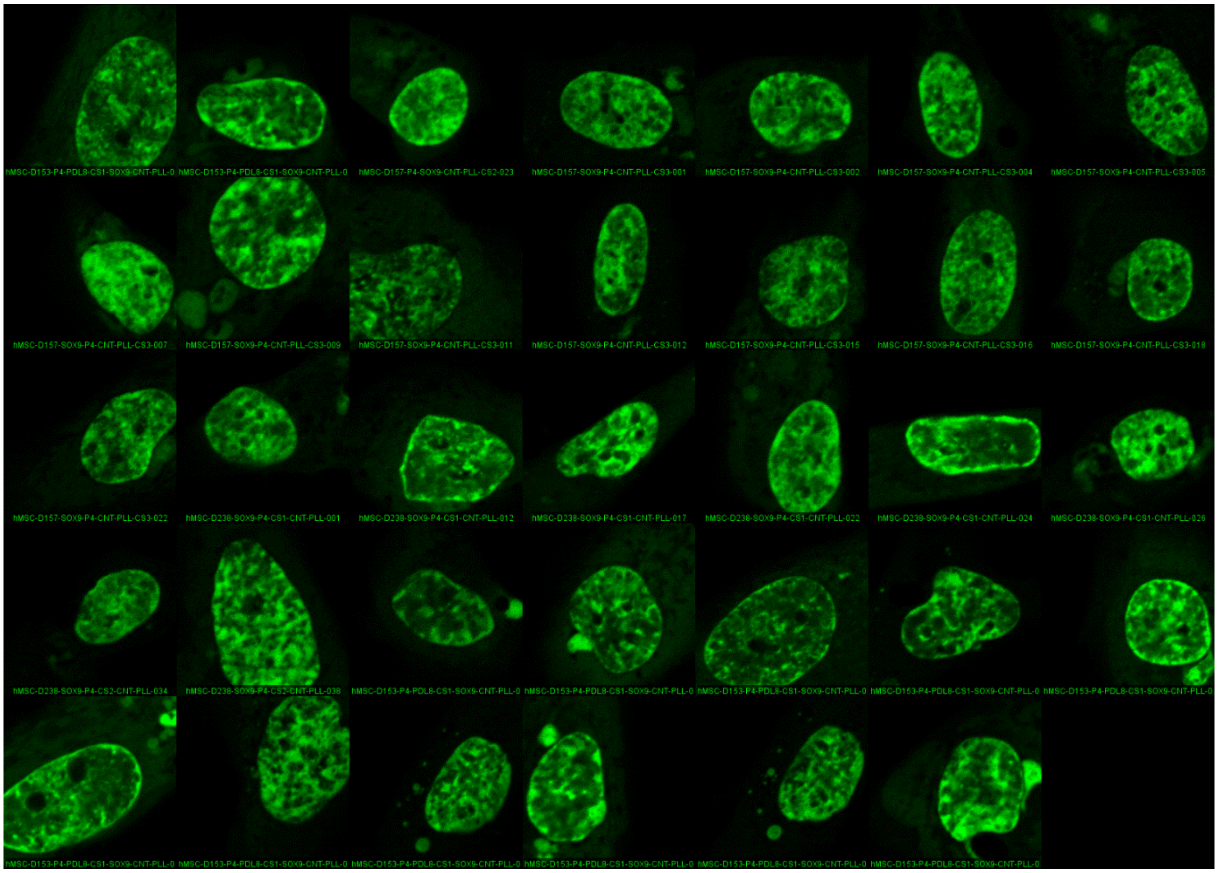


***Figure S10: Montage of nuclei showing nuclear localization pattern of SOX9-mGFP in the cluster A of control group (undifferentiated hMSCs).***
