## Supplementary material for "Mapping SOX9 transcriptional dynamics during multi-lineage differentiation of human mesenchymal stem cells": Figure S12

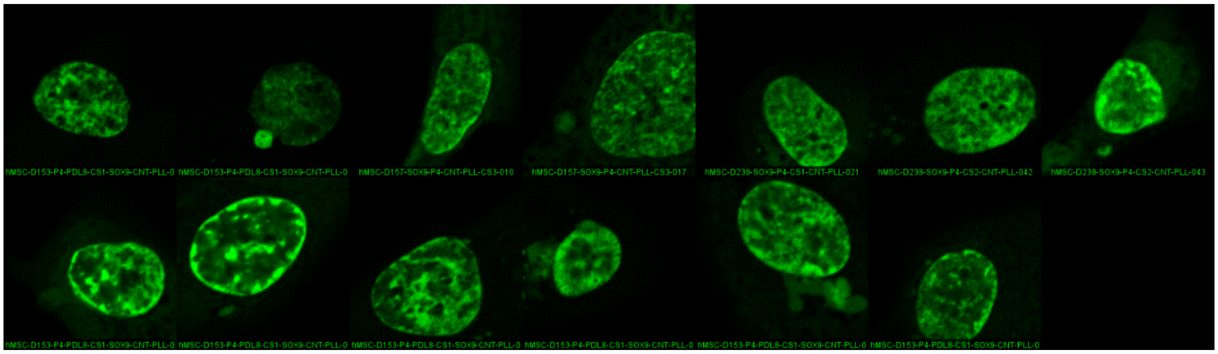


***Figure S12: Montage of nuclei showing nuclear localization pattern of SOX9-mGFP in the cluster C of control group (undifferentiated hMSCs).***
