## Supplementary material for "Mapping SOX9 transcriptional dynamics during multi-lineage differentiation of human mesenchymal stem cells": Figure S13

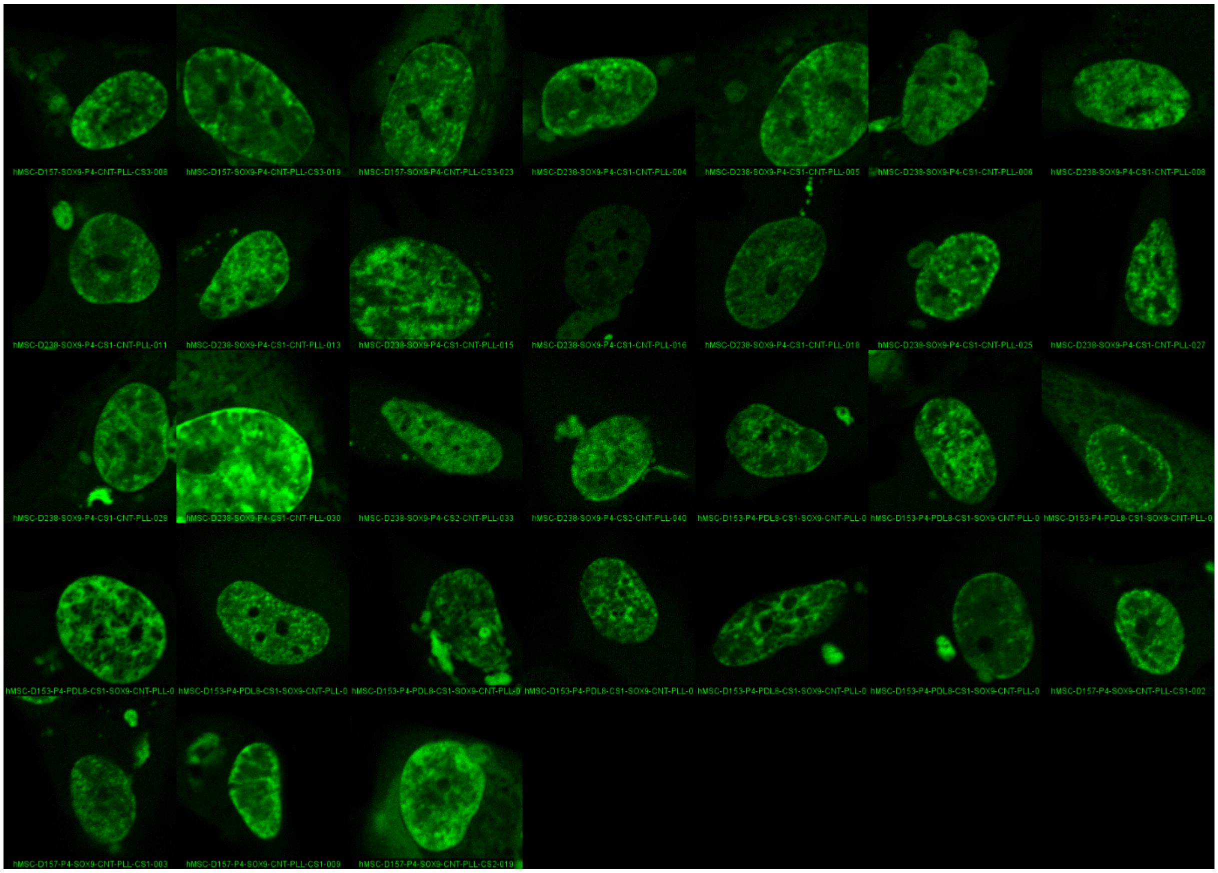


***Figure S13: Montage of nuclei showing nuclear localization pattern of SOX9-mGFP in the cluster D of control group (undifferentiated hMSCs).***
