## Supplementary material for "Mapping SOX9 transcriptional dynamics during multi-lineage differentiation of human mesenchymal stem cells": Figure S14

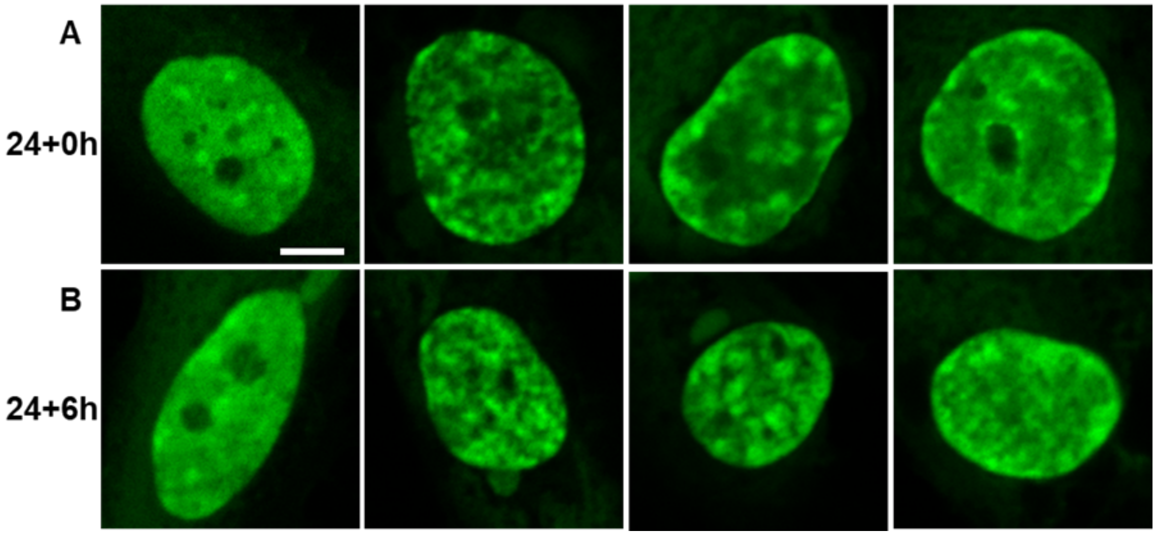


***Figure S14: Cell cycle synchronized hMSCs show at least four types of nuclear localization pattern of SOX9-mGFP.*** *(A) Cells were synchronized by maintaining them in the media without FBS for 24 hours. Cells were imaged at the end of 24 hours. (B) Cells synchronized for 24 hours were replaced with FBS containing media for 6 hours and cells were imaged at 30h (24 + 6h). This indicates that the different types of SOX9 nuclear localization pattern is not due to different stages of cell cycle. Scale bar: 5 µm.*
