## Supplementary material for "Mapping SOX9 transcriptional dynamics during multi-lineage differentiation of human mesenchymal stem cells": Figure S15

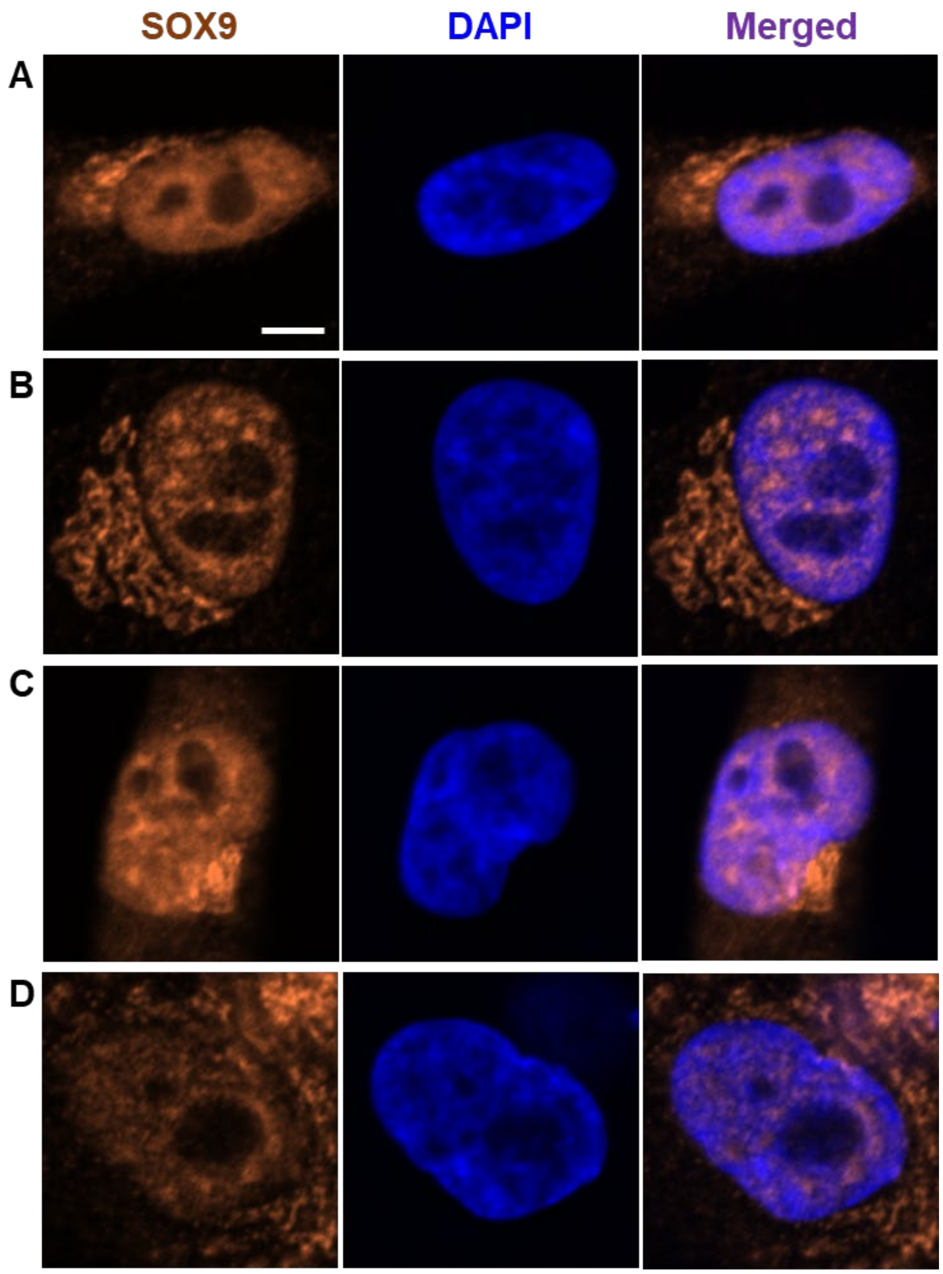


***Figure S15: Immunostaining of endogenous SOX9 in hMSCs show at least four types nuclear localization patterns in the undifferentiated hMSCs.*** This indicates that the differential nuclear localization patterns of SOX9 in the SOX9-mGFP transfected cells are not due to its overexpression. Scale bar: 5 µm.
