## Supplementary material for "Mapping SOX9 transcriptional dynamics during multi-lineage differentiation of human mesenchymal stem cells": Table S1

***Table S1: qPCR primer sequence.***

| Gene | Sequence |
| --- | --- |
| COL2A | 5’ CCAGATGACCTTCCTACGCC 3’ |
| | 5’ TTCAGGGCAGTGTACGTGAAC 3’ |
| ALPL | 5' ACAAGCACTCCCACTTCATC 3' |
| | 5' TTCAGCTCGTACTGCATGTC 3' |
| PPARγ | 5' GATGTCTCATAATGCCATCAGGTT 3' |
| | 5' GGATTCAGCTGGTCGATATCACT 3' |
| SOX9 | 5’ TGGGCAAGCTCTGGAGACTTC 3’ |
| | 5’ ATCCGGGTGGTCCTTCTTGTG 3’ |
| RPL13 | 5’ AAAAAGCGGATGGTGGTTC 3’ |
| | 5’ CTTCCGGTAGTGGATCTTGG 3’ |
