## Supplementary material for "Mapping SOX9 transcriptional dynamics during multi-lineage differentiation of human mesenchymal stem cells": Table S2

***Table S2: FRAP rates of SOX9-mGFP, number and percentage of cells in the individual clusters of undifferentiated hMSCs.***

| | **Immobile Fraction (%)** | **T-half of $A_2$ (sec)** | **T-half of $A_1$ (sec)** | **No. of cells** | **Percentage of cells** |
|---|---|---|---|---|---|
| Cluster A | 54.8 ± 4.4 | 12.56 ± 1.89 | 1.79 ± 0.29 | 34 | 26 |
| Cluster B | 53.2 ± 6.1 | 8.41 ± 2.04 | 1.25 ± 0.26 | 53 | 40 |
| Cluster C | 48.7 ± 4.8 | 18.29 ± 2.86 | 2.46 ± 0.63 | 13 | 10 |
| Cluster D | 38.5 ± 4.6 | 8.90 ± 2.44 | 1.54 ± 0.52 | 31 | 24 |
