## Supplementary material for "Mapping SOX9 transcriptional dynamics during multi-lineage differentiation of human mesenchymal stem cells": Table S3

***Table S3: FRAP rates of SOX9-mGFP, number and percentage of cells in the individual clusters of chondrogenically differentiating hMSCs. If same cluster is found more than two times, * indicates the significance pair.***

| | | **Immobile Fraction (%)** | **T-half of $A_2$ (sec)** | **T-half of $A_1$ (sec)** | **No. of cells** | **% of cells** | **P-Value** | | |
|---|---|---|---|---|---|---|---|---|---|
| | | | | | | | **IF** | **$A_2$** | **$A_1$** |
| Day 2 | Cluster E | 57.1 ± 8.3 | 11.47 ± 2.03 | 1.86 ± 0.33 | 18 | 14 | | | |
| | Cluster F | 53.8 ± 4.8 | 14.17 ± 5.14 | 2.36 ± 0.96 | 12 | 9 | | | |
| | Cluster G | 49.7 ± 8.6 | 10.37 ± 2.39 | 1.53 ± 0.33 | 53 | 41 | | | |
| | Cluster H | 42.5 ± 7.2 | 14.27 ± 4.78 | 1.94 ± 0.42 | 11 | 9 | | | |
| | Cluster I | 41.0 ± 8.7 | 8.97 ± 4.97 | 1.27 ± 0.43 | 14 | 11 | | | |
| | Cluster D | 35.9 ± 9.9 | 10.27 ± 2.25 | 1.49 ± 0.25 | 21 | 16 | 0.12 | 0.03 | 0.74 |
| Day 8 | Cluster A | 57.1 ± 4.1 | 11.32 ± 2.44 | 1.60 ± 0.32 | 15 | 11 | 0.11 | 0.13 | 0.10 |
| | Cluster J | 44.9 ± 9.3 | 20.26 ± 4.43 | 2.78 ± 0.32 | 13 | 10 | | | |
| | Cluster H* | 40.1 ± 5.3 | 15.35 ± 2.34 | 2.19 ± 0.28 | 21 | 15 | 0.33 | 0.48 | 0.11 |
| | Cluster D | 39.0 ± 6.0 | 9.30 ± 2.29 | 1.36 ± 0.29 | 64 | 47 | 0.53 | 0.49 | 0.30 |
| | Cluster K | 29.1 ± 6.4 | 11.16 ± 2.14 | 1.71 ± 0.31 | 23 | 17 | | | |
| Day 15 | Cluster F | 53.7 ± 4.3 | 19.73 ± 3.70 | 2.89 ± 0.33 | 8 | 6 | 0.51 | 0.02 | 0.56 |
| | Cluster L | 48.5 ± 2.6 | 11.63 ± 2.12 | 1.79 ± 0.22 | 17 | 13 | | | |
| | Cluster M | 39.3 ± 5.7 | 29.47 ± 3.75 | 3.89 ± 0.48 | 5 | 4 | | | |
| | Cluster D | 39.0 ± 4.2 | 9.17 ± 2.04 | 1.44 ± 0.36 | 52 | 39 | 0.64 | 0.57 | 0.63 |
| | Cluster H* | 36.9 ± 6.8 | 15.72 ± 2.37 | 2.32 ± 0.37 | 13 | 10 | 0.21 | 0.60 | 0.33 |
| | Cluster N | 25.5 ± 4.6 | 6.94 ± 2.56 | 1.01 ± 0.36 | 37 | 28 | | | |
