## Supplementary material for "Mapping SOX9 transcriptional dynamics during multi-lineage differentiation of human mesenchymal stem cells": Table S4

***Table S4: FRAP rates of SOX9-mGFP, number and percentage of cells in the individual clusters of osteogenically differentiating hMSCs. If same cluster is found more than two times, * and # indicate the significance pair.***

| | | Immobile Fraction (%) | T-half of $A_2$ (sec) | T-half of $A_1$ (sec) | No. of cells | % of cells | P-Value | | |
|---|---|---|---|---|---|---|---|---|---|
| | | | | | | | IF | $A_2$ | $A_1$ |
| Day 2 | Cluster E | 49.2 ± 6.2 | 6.92 ± 1.48 | 1.03 ± 0.16 | 18 | 15 | | | |
| | Cluster F# | 46.9 ± 5.5 | 10.98 ± 1.80 | 1.74 ± 0.27 | 53 | 43 | 0.20 | 0.02 | 0.09 |
| | Cluster C | 45.2 ± 8.9 | 17.66 ± 3.86 | 2.64 ± 0.27 | 17 | 14 | 0.45 | 0.23 | 0.16 |
| | Cluster G | 33.3 ± 4.0 | 7.06 ± 1.40 | 1.18 ± 0.20 | 9 | 7 | | | |
| | Cluster H | 31.0 ± 5.4 | 13.21 ± 3.37 | 1.80 ± 0.32 | 27 | 22 | | | |
| Day 8 | Cluster I | 58.4 ± 5.2 | 9.62 ± 1.60 | 1.58 ± 0.31 | 38 | 25 | | | |
| | Cluster E | 48.0 ± 3.9 | 7.20 ± 1.77 | 1.17 ± 0.28 | 36 | 23 | 0.88 | 0.56 | 0.05 |
| | Cluster C | 47.6 ± 10.1 | 19.87 ± 3.12 | 2.96 ± 0.49 | 12 | 8 | 0.94 | 0.20 | 0.04 |
| | Cluster F* | 46.1 ± 5.2 | 13.17 ± 2.52 | 1.97 ± 0.28 | 35 | 23 | 0.17 | 0.06 | 0.13 |
| | Cluster D | 37.2 ± 5.4 | 9.09 ± 2.30 | 1.39 ± 0.33 | 33 | 21 | 0.53 | 0.73 | 0.47 |
| Day 23 | Cluster F*# | 48.3 ± 5.8 | 12.09 ± 2.22 | 1.86 ± 0.35 | 45 | 32 | | | |
| | Cluster C | 46.8 ± 4.1 | 19.58 ± 1.65 | 3.18 ± 0.52 | 4 | 3 | 0.40 | 0.28 | 0.10 |
| | Cluster J | 42.9 ± 4.8 | 6.95 ± 1.99 | 1.06 ± 0.31 | 35 | 25 | | | |
| | Cluster K | 30.8 ± 4.1 | 9.09 ± 2.20 | 1.39 ± 0.39 | 41 | 29 | | | |
| | Cluster L | 30.6 ± 4.9 | 15.93 ± 3.89 | 2.24 ± 0.29 | 16 | 11 | | | |
