## Supplementary material for "Mapping SOX9 transcriptional dynamics during multi-lineage differentiation of human mesenchymal stem cells": Table S5

***Table S5: FRAP rates of SOX9-mGFP, number and percentage of cells in the individual clusters of adipogenically differentiating hMSCs. If same cluster is found more than two times, * and # indicate the significance pair.***

| | | Immobile Fraction (%) | T-half of $A_2$ (sec) | T-half of $A_1$ (sec) | No. of cells | % of cells | P-Value | | |
|---|---|---|---|---|---|---|---|---|---|
| | | | | | | | IF | $A_2$ | $A_1$ |
| Day 2 | Cluster E* | 57.7 ± 8.4 | 16.43 ± 2.44 | 2.31 ± 0.56 | 15 | 12 | 0.08 | 0.06 | 0.79 |
| | Cluster B | 53.7 ± 5.7 | 8.89 ± 1.52 | 1.38 ± 0.25 | 44 | 34 | 0.79 | 0.30 | 0.03 |
| | Cluster F | 41.5 ± 6.8 | 12.77 ± 2.27 | 1.85 ± 0.35 | 38 | 29 | | | |
| | Cluster G | 37.4 ± 6.7 | 6.47 ± 1.92 | 0.98 ± 0.26 | 32 | 25 | | | |
| Day 8 | Cluster H | 55.2 ± 5.6 | 10.99 ± 1.58 | 1.58 ± 0.20 | 36 | 29 | | | |
| | Cluster E*# | 54.6 ± 5.8 | 14.89 ± 1.77 | 2.26 ± 0.29 | 22 | 18 | | | |
| | Cluster I | 51.0 ± 6.6 | 7.10 ± 2.00 | 1.07 ± 0.30 | 33 | 26 | | | |
| | Cluster J | 36.5 ± 6.2 | 15.14 ± 4.30 | 2.17 ± 0.17 | 11 | 9 | | | |
| | Cluster K | 32.1 ± 4.6 | 6.76 ± 2.06 | 1.06 ± 0.35 | 23 | 18 | | | |
| Day 23 | Cluster E# | 57.0 ± 4.3 | 14.96 ± 2.28 | 2.13 ± 0.29 | 17 | 13 | 0.19 | 0.92 | 0.18 |
| | Cluster H | 52.7 ± 5.4 | 10.53 ± 1.37 | 1.60 ± 0.21 | 38 | 30 | 0.05 | 0.19 | 0.69 |
| | Cluster L | 46.8 ± 6.3 | 7.19 ± 1.53 | 1.04 ± 0.19 | 42 | 33 | | | |
| | Cluster D | 36.4 ± 4.6 | 10.40 ± 2.08 | 1.51 ± 0.25 | 15 | 12 | 0.23 | 0.06 | 0.69 |
| | Cluster J | 32.3 ± 7.9 | 13.96 ± 1.90 | 2.45 ± 0.33 | 8 | 6 | 0.27 | 0.65 | 0.04 |
| | Cluster M | 30.1 ± 3.4 | 4.51 ± 1.11 | 0.71 ± 0.17 | 8 | 6 | | | |
