## Supplementary material for "Mapping SOX9 transcriptional dynamics during multi-lineage differentiation of human mesenchymal stem cells": Table S6

*Table S6: FRAP rates of SOX9-mGFP in osteogenically differentiating hMSCs at day 23, with and without FACS sorting for CD10+ and/or CD92+ cells. *significance pair.*

| | | Immobile Fraction (%) | T-half of $A_2$ (sec) | T-half of $A_1$ (sec) | P-Value | | |
|---|---|---|---|---|---|---|---|
| | | | | | IF | $A_2$ | $A_1$ |
| FACS Sorted | CD10+* | 39.0 ± 8.2 | 8.55 ± 2.67 | 1.38 ± 0.40 | | | |
| | CD92+ | 36.1 ± 8.9 | 11.25 ± 2.96 | 1.58 ± 0.39 | | | |
| | CD10+92+ | 43.5 ± 6.4 | 11.99 ± 3.12 | 1.63 ± 0.35 | | | |
| Day 23 | Cluster F | 48.3 ± 5.8 | 12.09 ± 2.22 | 1.86 ± 0.35 | | | |
| | Cluster C | 46.8 ± 4.1 | 19.58 ± 1.65 | 3.18 ± 0.52 | | | |
| | Cluster J | 42.9 ± 4.8 | 6.95 ± 1.99 | 1.06 ± 0.31 | | | |
| | Cluster K* | 30.8 ± 4.1 | 9.09 ± 2.20 | 1.39 ± 0.39 | 0.0001 | 0.29 | 0.67 |
| | Cluster L | 30.6 ± 4.9 | 15.93 ± 3.89 | 2.24 ± 0.29 | | | |
