## Supplementary material for "Mapping SOX9 transcriptional dynamics during multi-lineage differentiation of human mesenchymal stem cells": Table S7

***Table S7: Comparing FRAP rates of SOX9-mGFP among undifferentiated hMSCs (control), chondrogenically differentiated hMSCs (day 15), healthy and OA hPCs.***

| | | Immobile Fraction (%) | T-half of $A_2$ (sec) | T-half of $A_1$ (sec) | P-Value | | |
|---|---|---|---|---|---|---|---|
| | | | | | IF | $A_2$ | $A_1$ |
| hMSC Control | Cluster A* | 54.8 ± 4.4 | 12.56 ± 1.89 | 1.79 ± 0.29 | | | |
| | Cluster B | 53.2 ± 6.1 | 8.41 ± 2.04 | 1.25 ± 0.26 | | | |
| | Cluster C | 48.7 ± 4.8 | 18.29 ± 2.86 | 2.46 ± 0.63 | | | |
| | Cluster D | 38.5 ± 4.6 | 8.90 ± 2.44 | 1.54 ± 0.52 | | | |
| hMSC CD Day 15 | Cluster F | 53.7 ± 4.3 | 19.73 ± 3.70 | 2.89 ± 0.33 | | | |
| | Cluster L | 48.5 ± 2.6 | 11.63 ± 2.12 | 1.79 ± 0.22 | | | |
| | Cluster M | 39.3 ± 5.7 | 29.47 ± 3.75 | 3.89 ± 0.48 | | | |
| | Cluster D | 39.0 ± 4.2 | 9.17 ± 2.04 | 1.44 ± 0.36 | | | |
| | Cluster H | 36.9 ± 6.8 | 15.72 ± 2.37 | 2.32 ± 0.37 | | | |
| | Cluster N | 25.5 ± 4.6 | 6.94 ± 2.56 | 1.01 ± 0.36 | | | |
| hPCs | HL Cluster 1 | 59.3 ± 3.3 | 14.05 ± 1.97 | 1.98 ± 0.36 | | | |
| | HL Cluster 2* | 68.0 ± 4.4 | 10.84 ± 3.46 | 1.63 ± 0.47 | 0.008 | 0.09 | 0.14 |
| | OA Cluster 1 | 47.4 ± 4.6 | 7.25 ± 2.48 | 1.07 ± 0.38 | | | |
| | OA Cluster 2 | 58.4 ± 3.4 | 9.38 ± 2.68 | 1.24 ± 0.29 | | | |
