## Supplementary material for "Mapping SOX9 transcriptional dynamics during multi-lineage differentiation of human mesenchymal stem cells": Figure S1

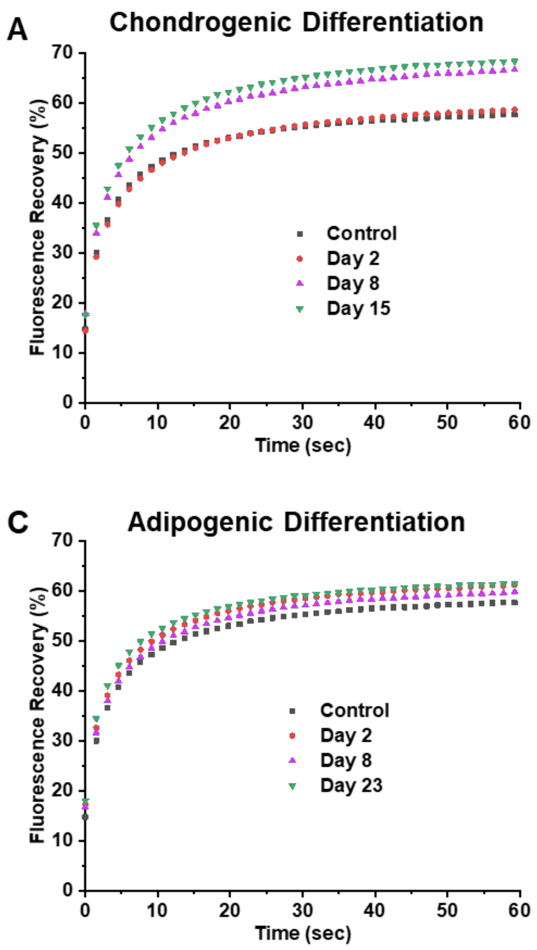


***Figure S1: Averaged FRAP curves of SOX9-mGFP (without clustering), per time point, during (A) chondrogenic, (B) osteogenic and (C) adipogenic differentiation of hMSCs show that SOX9 mobility only increase during any differentiation lineage as compared to undifferentiated control.*** FRAP measurements were performed in three donors separately and combined for analysis (n = ≥ 125 cells, per time point, three donors together).
