## Supplementary material for "Mapping SOX9 transcriptional dynamics during multi-lineage differentiation of human mesenchymal stem cells": Figure S3

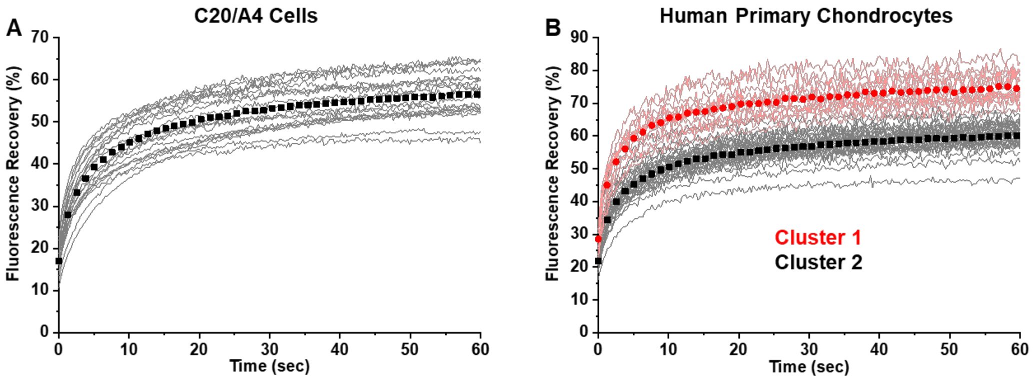


*Figure S3: FRAP cures of SOX9-mGFP in C20/A4 cells (A) show only one type of mobility pattern and human primary chondrocytes isolated from OA joint (B) show two clusters of mobility pattern.*
