## Supplementary material for "Mapping SOX9 transcriptional dynamics during multi-lineage differentiation of human mesenchymal stem cells": Figure S5

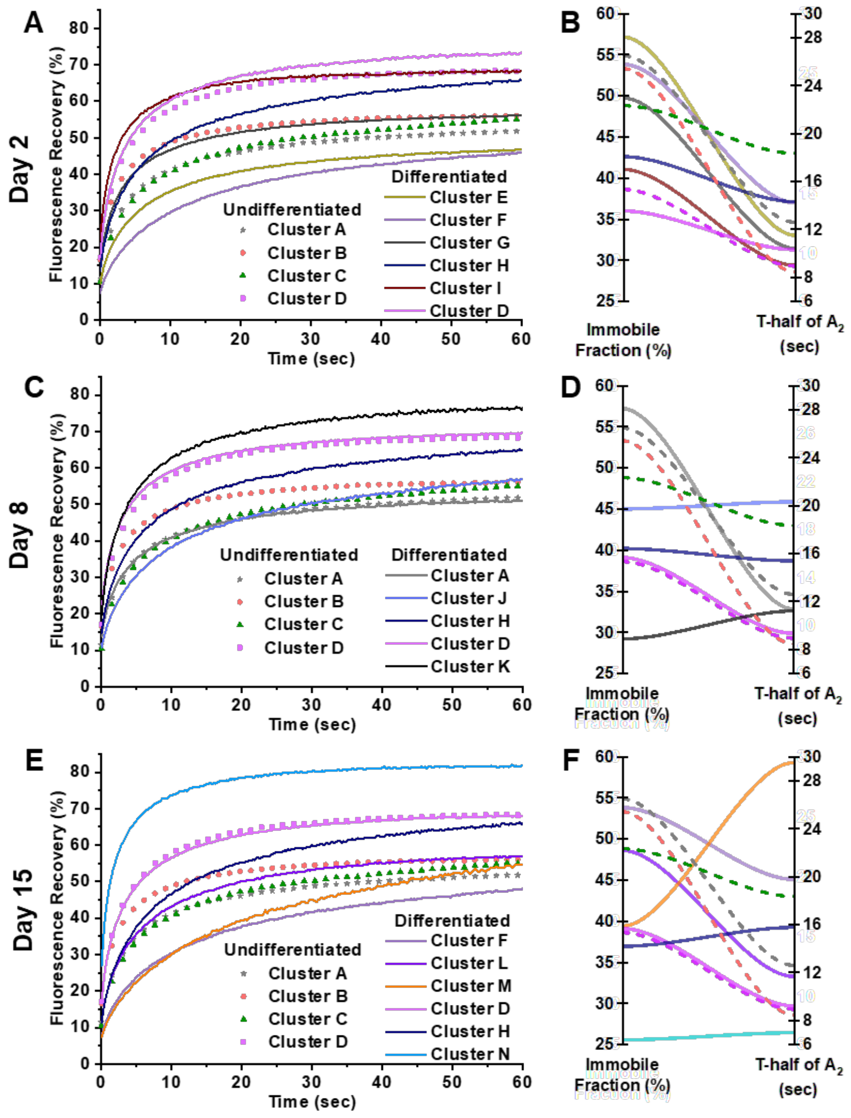


***Figure S5: Mobility of SOX9-mGFP in undifferentiated (symbols or dash line) and chondrogenically differentiating (continuous line) hMSCs as measured by FRAP.*** *Unsupervised hierarchical clustering identified at least four distinct clusters in undifferentiated control group based on the FRAP variables IF, t½ of $A_1$ and $A_2$. In the differentiated group, six clusters with distinct dynamic rates appeared at day 2 (A and B), 15 (E and F) and five clusters appeared at day 8 (C and D). FRAP recovery curves on the left show changes in the mobility pattern of SOX9-mGFP in the chondrogenically differentiating clusters as compared to clusters in undifferentiated cells. Parallel plot on the right show the relationship between IF and t½ of $A_2$ and its changes between undifferentiated (dashed line) and differentiating hMSCs (continuous line). Three hMSC donors were used for this study. Combined data from all the donors are shown here. n ≥ 129, per time point.*
